# Transcriptomic analysis of the human habenula in schizophrenia

**DOI:** 10.1101/2024.02.26.582081

**Authors:** Ege A. Yalcinbas, Bukola Ajanaku, Erik D. Nelson, Renee Garcia-Flores, Nicholas J. Eagles, Kelsey D. Montgomery, Joshua M. Stolz, Joshua Wu, Heena R. Divecha, Atharv Chandra, Rahul A. Bharadwaj, Svitlana Bach, Anandita Rajpurohit, Ran Tao, Geo Pertea, Joo-Heon Shin, Joel E. Kleinman, Thomas M. Hyde, Daniel R. Weinberger, Louise A. Huuki-Myers, Leonardo Collado-Torres, Kristen R. Maynard

## Abstract

Pathophysiology of many neuropsychiatric disorders, including schizophrenia (SCZD), is linked to habenula (Hb) function. While pharmacotherapies and deep brain stimulation targeting the Hb are emerging as promising therapeutic treatments, little is known about the cell type-specific transcriptomic organization of the human Hb or how it is altered in SCZD. Here we define the molecular neuroanatomy of the human Hb and identify transcriptomic changes in individuals with SCZD compared to neurotypical controls. Utilizing Hb-enriched postmortem human brain tissue, we performed single nucleus RNA-sequencing (snRNA-seq; n=7 neurotypical donors) and identified 17 molecularly defined Hb cell types across 16,437 nuclei, including 3 medial and 7 lateral Hb populations, several of which were conserved between rodents and humans. Single molecule fluorescent *in situ* hybridization (smFISH; n=3 neurotypical donors) validated snRNA-seq Hb cell types and mapped their spatial locations. Bulk RNA-sequencing and cell type deconvolution in Hb-enriched tissue from 35 individuals with SCZD and 33 neurotypical controls yielded 45 SCZD-associated differentially expressed genes (DEGs, FDR < 0.05), with 32 (71%) unique to Hb-enriched tissue. eQTL analysis identified 717 independent SNP-gene pairs (FDR < 0.05), where either the SNP is a SCZD risk variant (16 pairs) or the gene is a SCZD DEG (7 pairs). eQTL and SCZD risk colocalization analysis identified 16 colocalized genes. These results identify topographically organized cell types with distinct molecular signatures in the human Hb and demonstrate unique genetic changes associated with SCZD, thereby providing novel molecular insights into the role of Hb in neuropsychiatric disorders.

**One Sentence Summary:** Transcriptomic analysis of the human habenula and identification of molecular changes associated with schizophrenia risk and illness state.

## Introduction

The habenula (Hb) is a small epithalamic brain structure implicated in several neuropsychiatric conditions, including mood and anxiety disorders, substance use disorders, and schizophrenia (SCZD) *(1–3)*. Anatomically, it is divided into molecularly and functionally distinct medial (MHb) and lateral (LHb) subdivisions with discrete connections to monoaminergic hubs, including dopamine, serotonin, and norepinephrine centers *(2)*. As a circuit hub involved in motivated behaviors and affective states, the Hb has emerged as a promising therapeutic target *(4–9)*. Imaging studies in SCZD patients have sought to identify aberrations in Hb anatomy, connectivity, and activity, but challenges in segmenting this small midline structure have hindered investigation into its subregions and limited conclusive findings *(10–14)*. Similarly, there is a paucity of molecular studies in postmortem human Hb due to its challenging size and location *(15–18)*. While bulk RNA-sequencing (RNA-seq) studies identified gene expression changes associated with SCZD in other brain regions *(19–21)*, the molecular landscape of the Hb in SCZD has not yet been investigated.

Single cell RNA-sequencing studies in animal models found diverse cell populations with unique molecular signatures across LHb and MHb *(22–24)*. However, it is unclear whether these Hb cell types are conserved in the human brain. This information is important for understanding biological vulnerabilities underlying human disease, which is critical for improving translational research outcomes *(25)*. Given the central role of the Hb in modulating dopaminergic, serotonergic, and cholinergic pathways, molecular profiling in this region can reveal novel mechanistic insights into disease pathogenesis, especially regarding the nuanced role of dopamine dysregulation in SCZD. For example, the fact that many antipsychotic medications act through dopamine and serotonin receptors warrants a deeper exploration of Hb cell types and signaling pathways contributing to downstream monoaminergic dysfunction*(26)*. Additionally, SCZD is associated with nicotine dependence, and smoking is reported to improve cognitive symptoms *(27–29)*. The MHb is enriched in cholinergic populations implicated in nicotine addiction *(30–32)*, but a link between cholinergic signaling, nicotine dependence, and psychotic disorders as it relates to the human Hb has yet to be elucidated *(33–36)*.

To address these gaps in knowledge, we generated the first single cell transcriptomic atlas of the healthy human Hb and evaluated the topography and cross-species convergence of MHb and LHb cell types. To understand molecular changes in the human Hb associated with SCZD, we performed bulk RNA-seq and expression quantitative trait locus (eQTL) analysis in Hb-enriched tissue from SCZD and neurotypical control donors. We identified gene expression changes and eQTLs unique to Hb-enriched tissue compared to other brain regions in the context of SCZD. Our findings support previous hypotheses of SCZD etiology that implicate neurodevelopmental processes *(37–40)*, while providing additional molecular insights into Hb cell types that may contribute to SCZD pathogenesis.

## Results

### Identification of distinct human Hb cell types

To generate a molecular atlas of human Hb cell populations, we performed snRNA-seq on neuronal (NeuN+) and non-neuronal nuclei from Hb-enriched tissue from 7 neurotypical control donors (**Fig. 1A; Table S1**). Following quality control, 16,437 single nucleus transcriptomes were included in downstream analyses and annotated at broad resolution based on canonical marker gene expression (**Fig. S1-5; Table S2**). Graph-based clustering identified 17 molecularly defined cell types, including non-neuronal populations (oligodendrocytes, astrocytes, microglia, oligodendrocyte precursor cells, and endothelial cells; n=4,101 nuclei), thalamic neurons (excitatory and inhibitory; n=9,412 nuclei), and Hb neurons (7 lateral [L] and 3 medial [M] Hb subpopulations; n=2,924 nuclei) (**Fig. 1B,C**; **Fig. S13-13; Supplemental Methods**). Principal component analysis revealed that the second and third components of variation differentiated Hb from thalamic neurons, and neuronal from non-neuronal cell types, respectively (**Fig. S15**). The small size and midline location of the Hb resulted in variability in epithalamus dissections across donors; thus, our samples contained varying proportions of thalamic and Hb neurons, as well as neuronal versus non-neuronal cell types (**Fig. 1D**). However, neuronal enrichment with NeuN+ sorting in a subset of samples increased the capture rate of Hb neurons, and we observed the presence of LHb and MHb neuronal subpopulations in most samples.

**Fig. 1.**
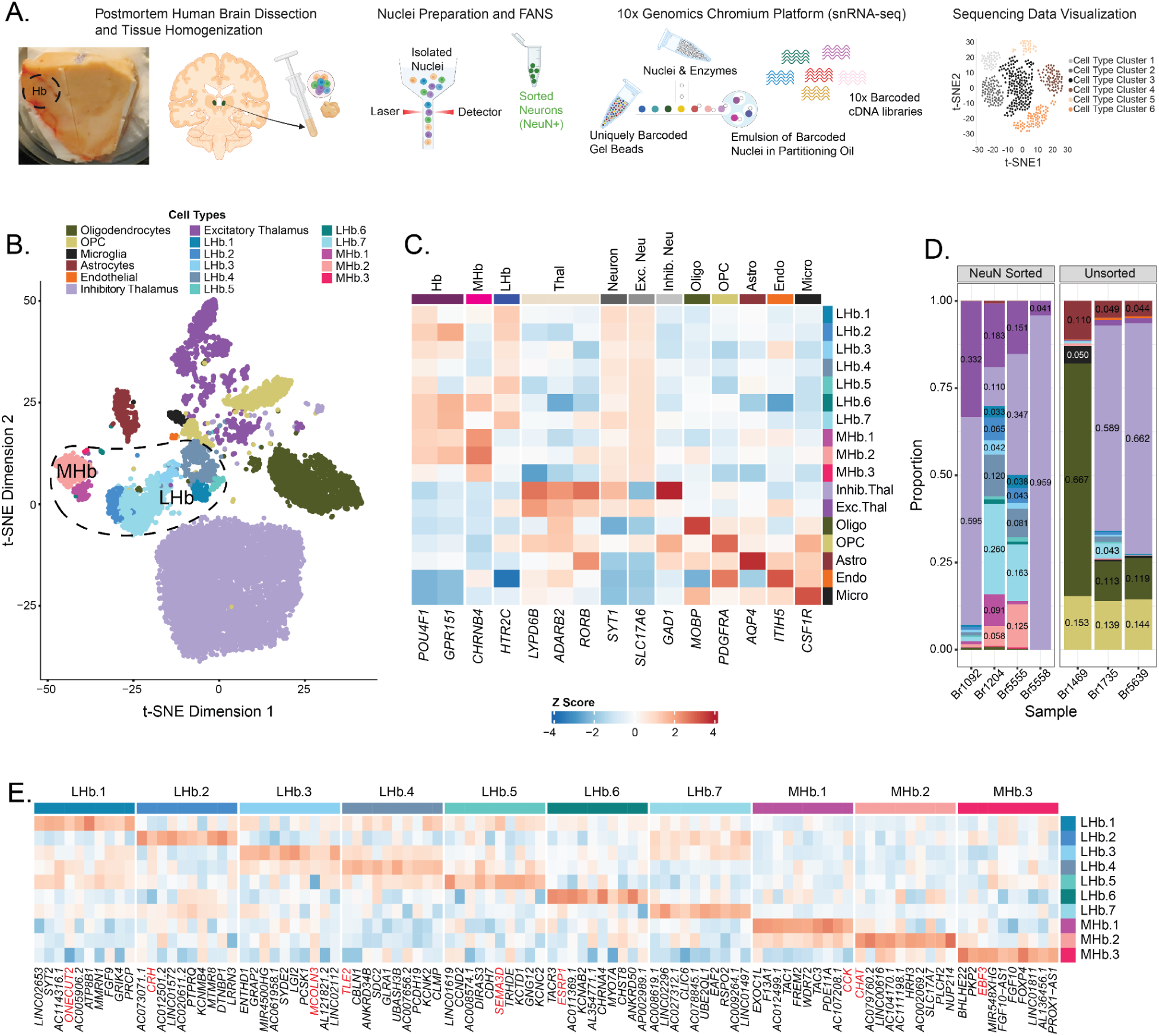
Single cell transcriptomic atlas of the human habenula (Hb). **A)** Overview of nuclei preparation and single nucleus RNA-sequencing (snRNA-seq) data generation using the 10x Genomics Chromium Single Cell Gene Expression platform **B)** t-SNE plot of 16,437 quality-controlled single nucleus transcriptomes (colored by cell type) from the epithalamus of 7 neurotypical control donors. Dashed circle highlights lateral (L) and medial (M) Hb cell type clusters (n=2,214 LHb and n=710 MHb nuclei). **C)** Heatmap depicting expression of known marker genes (x-axis) that were used to annotate broad cell types as well as habenula vs. thalamic neurons. The color of each square (blue to red) indicates z-scored relative expression of the corresponding marker gene. **D)** Proportion plot displaying cell type composition of each sample. NeuN+ sorted samples were enriched for neurons and therefore contain more neuronal cell types. **E)** Heatmap highlighting the top 10 mean ratio marker genes (x-axis) for each identified LHb and MHb neuronal subpopulation. The color of each square (blue to red) indicates z-scored relative expression of the corresponding marker gene. Marker genes highlighted in red text were further validated using single molecule fluorescent *in situ* hybridization (smFISH, see Fig. 3).

Broad-level cell labels were assigned based on previously established marker genes from human, rodent, and zebrafish studies (**Fig. 1C**) *(22–24, 41–44)*. As expected, *POU4F1* and *GPR151* were enriched in human Hb subpopulations compared to thalamic and non-neuronal populations *(17, 45–48)*. We also identified *CHRNB4* and *HTR2C* as marker genes for MHb and LHb subpopulations, respectively (**Table S6**). *CHRNB4* encodes a nicotinic acetylcholine receptor (nAChR) subunit, and is an MHb marker gene in both rodents and humans along with *CHRNA3*, another nAChR subunit listed as a top mean ratio marker gene for MHb *(23)*. These two genes are in a conserved gene locus implicated in nicotine dependence *(49, 50)*. *HTR2C,* which encodes a G-protein-coupled receptor (GPCR) serotonin receptor, is also a rodent LHb marker gene *(22, 23)*. To better understand cell type diversity within anatomical subdivisions of Hb, we identified top mean ratio marker genes for the 10 LHb and MHb subpopulations (**Fig. 1E; Table S3; Supplemental Methods**). This allowed us to gain functional insights into different LHb and MHb subpopulations, such as the identification of cholinergic neurons (MHb.2) expressing high levels of *CHAT* and the identification of a LHb.2 subpopulation expressing high levels of *CRH* and *OPRM1*.

### Conservation of Hb cell types from preclinical rodent model in human Hb

To evaluate whether Hb cell types found in rodents are conserved in human Hb, we conducted a cross-species correlation analysis with mouse Hb single cell RNA-seq data *(22)*. We observed positive correlations between the human and mouse datasets across broad cell type categories, including astrocytes, microglia, oligodendrocytes, and neurons (**Fig. 2A**). A correlation analysis restricted to human Hb (3 MHb, 7 LHb) and mouse Hb neuronal cell types (6 MHb, 6 LHb) revealed some cross-species convergence of Hb neurons (**Fig. 2B**). Human MHb subpopulations were most strongly correlated with mouse MHb subpopulations, and the same relationship held true for the LHb subpopulations, with the exception of human LHb.6 which most strongly correlated with mouse MHb.1. Of particular interest was the convergence of human MHb.1 and MHb.2 with mouse MHb subpopulations. Human MHb.1 neurons express relatively high levels of *CCK*, *TAC1*, and *TAC3,* which encode various neuropeptides; human MHb.2 neurons are cholinergic as evidenced by relative high levels of *CHAT* expression (**Fig. 1E**). Neuronal subpopulations enriched in these marker genes were previously identified in rodent Hb *(22, 23)*. Among the LHb subpopulations, human LHb.4, LHb.3, and LHb.2 had the strongest positive correlation with mouse LHb.3, LHb.1, and LHb.5, respectively, suggesting cross-species convergence of these molecularly distinct neuronal cell types.

**Fig. 2.**
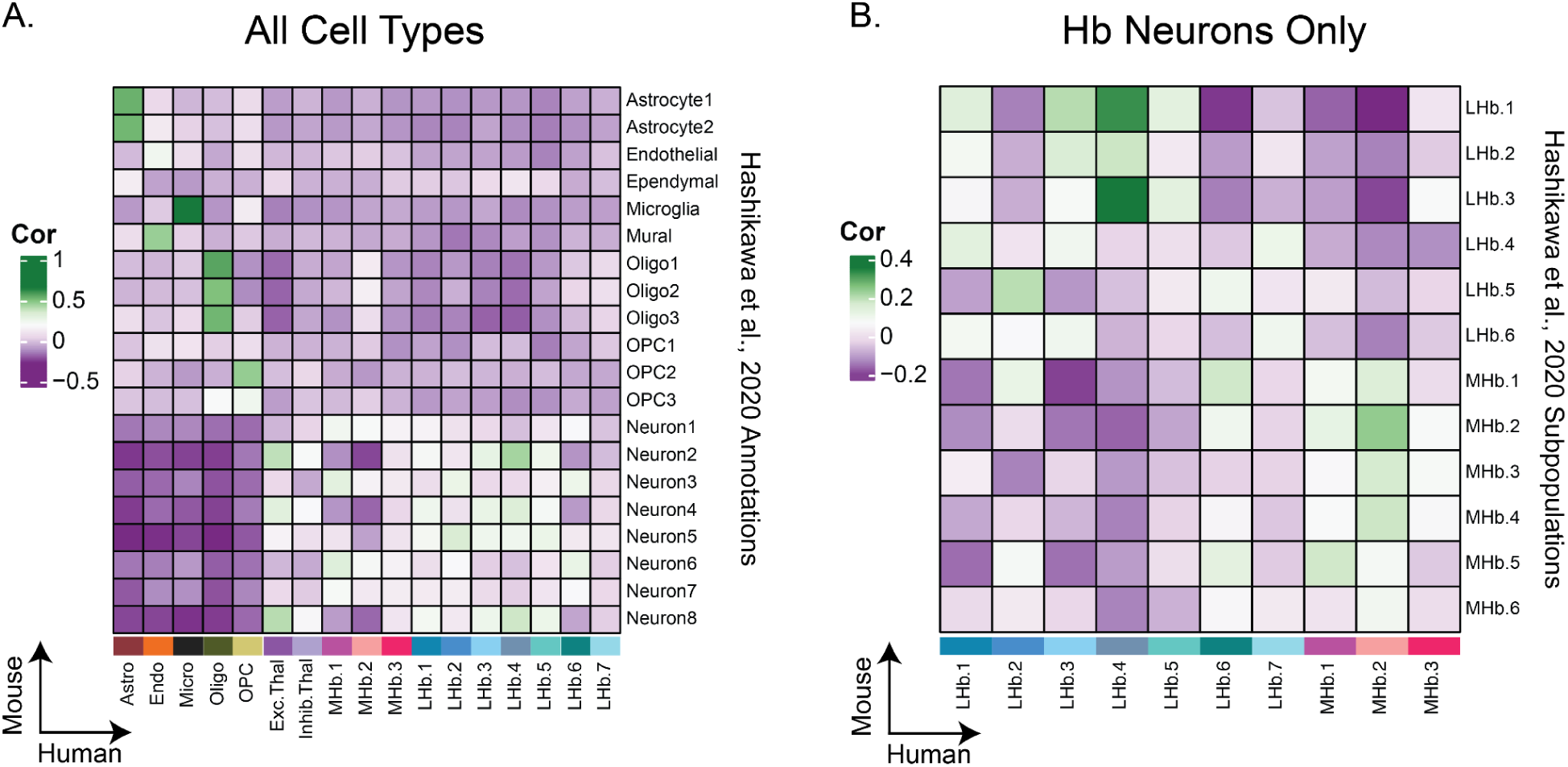
Conservation of LHb and MHb neuronal subpopulations from preclinical rodent model in human Hb. **A)** Pearson correlation heatmap of *t*-statistics obtained from a cell type differential gene expression (DGE) analysis conducted on a mouse Hb scRNA-seq dataset *(22)* along the y-axis, and *t*-statistics from a DGE analysis of our human cell types identified in Fig. 1B along the x-axis. Green squares show positive correlation, i.e. convergence of cell type-specific gene expression patterns between species. **B)** Correlation heatmap obtained similarly to A, but subset to Hb neuronal subpopulations.

### Topography of human Hb cell types

Given that rodent LHb and MHb neuronal subpopulations display topography *(23, 24)*, we validated the spatial organization of select human Hb subpopulations using single molecule fluorescent *in situ* hybridization (smFISH; n=2-3 donors per experiment; **Table S4**, **Table S5**). We used three probe combinations each targeting 4 mean ratio marker genes to visualize the following subpopulations (many of which could be identified by one marker gene, or a combination of 2 genes): LHb.1, LH.4, and LHb.5/1 (*ONECUT2*, *TLE2*, *SEMA3D*); LHb.2, LHb.3, and LHb.6 (*CRH*, *MCOLN3*, *ESRP1*); MHb.1, MHb.2, and MHb.3 (*CCK*, *CHAT*, *EBF3*) (**Fig. 3**). Across donors, we observed spatial biases in some human Hb cell type locations (**Fig. S6-8**). For example, cells that highly expressed *ESRP1* (putative LHb.6 neurons) displayed a dorsal bias (**Fig. 3B**). Furthermore, cells that highly expressed *CHAT* (putative MHb.2 neurons) clustered separately from other MHb cell types (**Fig. 3C**). Taken together, smFISH results validated snRNA-seq neuronal subpopulations and corroborated findings in mice that molecularly defined LHb and MHb subpopulations show distinct spatial organization.

**Fig. 3.**
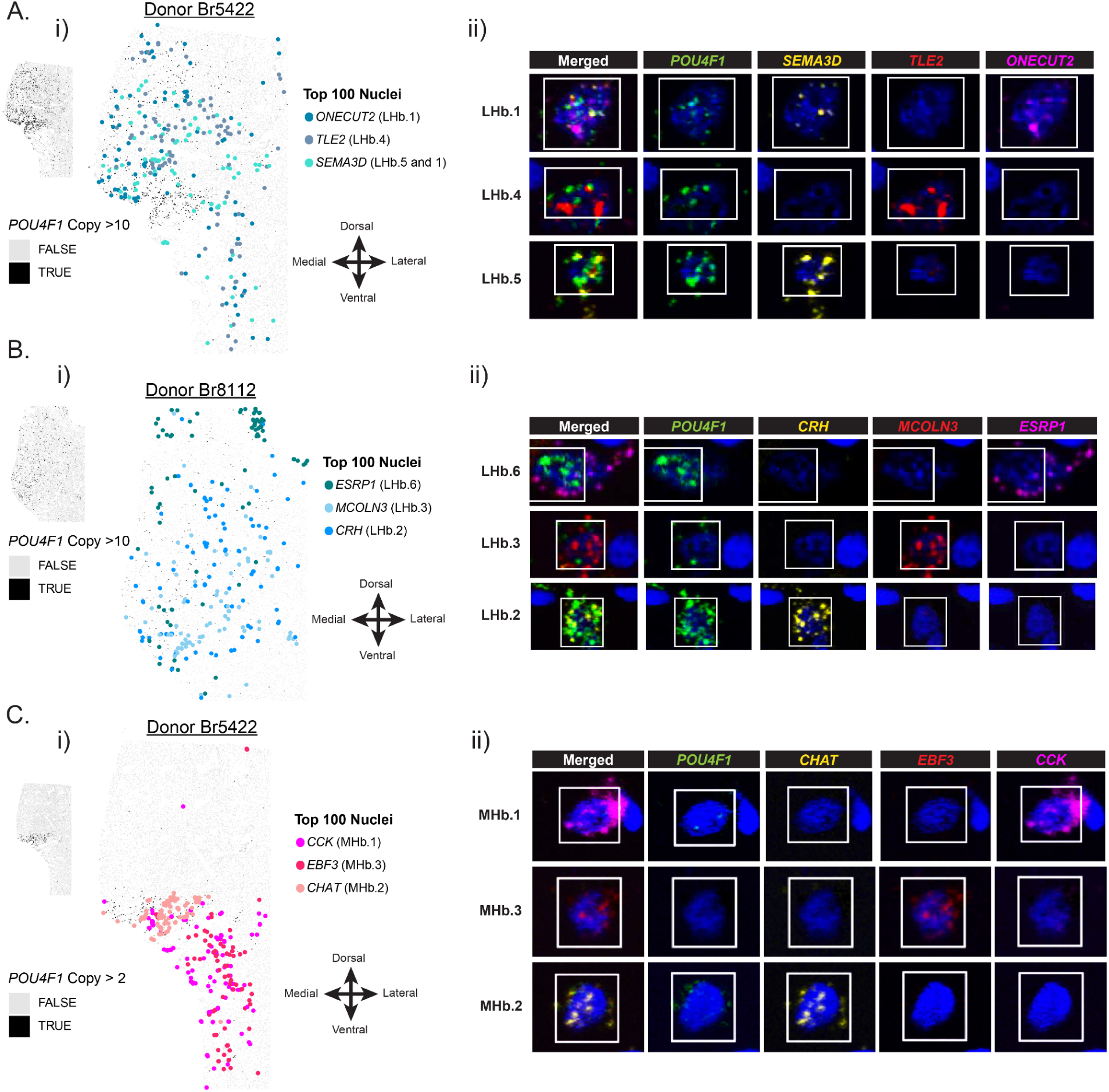
Spatial organization of molecularly-defined LHb and MHb neuronal subpopulations in postmortem human tissue. Spatial plots displaying the topographic arrangement of top cells that express the marker genes chosen for RNAScope validation of Hb subpopulations. **A**) (**i**) *ONECUT2, TLE2*, *SEMA3D* marker gene expression *in situ*, with each detected cell object in the tissue section plotted as a gray or black rectangle. Cells in black have > 10 transcript copies of the Hb-wide marker gene *POU4F1*, indicating the presence of habenula. Colored circles mark the spatial location of the top 100 cells that most robustly express each marker gene. If a cell was ranked in the top 100 for more than one marker gene (see confusion matrices in **Fig. S6B**), it was colored by the gene for which it had the most number of transcript copies. (**ii**) RNAScope images of example top ranked cells that highly express *ONECUT2, TLE2*, *SEMA3D* at high magnification. **B)** (**i**) Spatial plots as described in **A** for *ESRP1*, *MCOLN3*, *CRH* marker gene expression *in situ*. (**ii**) RNAScope images of example top ranked cells that highly express *ESRP1*, *MCOLN3*, *CRH* at high magnification. **C)** (**i**) Spatial plots as described in **A** for *CCK*, *EBF3*, *CHAT* marker gene expression in situ. (**ii**) RNAScope images of example top ranked cells that highly express *CCK*, *EBF3*, *CHAT* at high magnification.

### UNIQUE SCHIZOPHRENIA-ASSOCIATED TRANSCRIPTOMIC CHANGES IN HUMAN HB COMPARED TO OTHER BRAIN REGIONS

To investigate transcriptomic changes associated with SCZD diagnosis in the human Hb, we performed bulk RNA-seq of postmortem human Hb-enriched tissue samples from 35 SCZD cases and 34 neurotypical controls (**Fig. 4A**, **Table S1**, **Fig. S10**). Due to inclusion of surrounding thalamic tissue during epithalamus dissections, we leveraged our human Hb snRNA-seq dataset (**Fig. 1**) to implement cell type deconvolution of the bulk RNA-seq data. Samples contained some proportion of Hb cell types as estimated by utilizing the top 25 mean ratio marker genes for broad cell types (**Fig. S11**, **Fig. S12**, **Fig. S16**, **Table S6**). The one control sample that did not meet this criterion was dropped from further analyses.

**Fig. 4.**
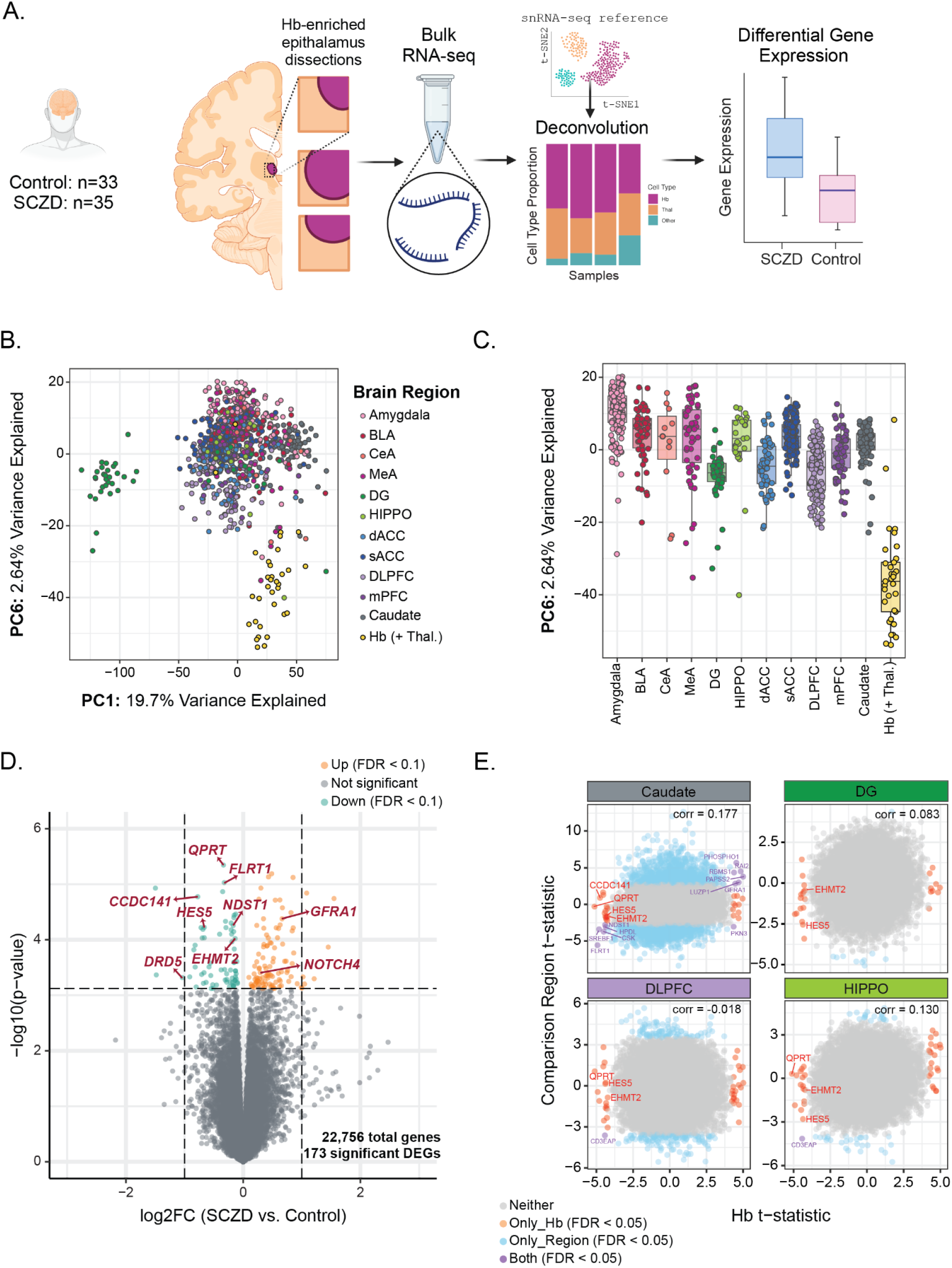
Identification of unique differentially expressed genes (DEGs) in Hb-enriched thalamus of Schizophrenia (SCZD) vs. Control cases. **A)** Study design and cell type deconvolution of postmortem human bulk RNA-seq samples using Hb single nucleus RNA-seq data (see Fig. 1 and **Fig. S12**). **B)** Principal component analysis of postmortem human bulk RNA-seq data from neurotypical control tissue samples of Hb-enriched thalamus [Hb (+Thal.)] and 11 other brain regions (combined n = 817): amygdala, basolateral amygdala (BLA), central amygdala (CeA), caudate, dorsal anterior cingulate cortex (dACC), dentate gyrus (DG), dorsolateral prefrontal cortex (DLPFC), hippocampus (HIPPO), medial amygdala (MeA), medial prefrontal cortex (mPFC), and subgenual anterior cingulate cortex (sACC). PC1 vs. PC6 scatter plot depicts PC6 separating Hb (+Thal.) from other regions. **C)** Boxplot of PC6 across 12 tested brain regions. **D)** Volcano plot of DGE analysis showing. upregulated genes (FDR < 0.1, orange points) and downregulated genes (FDR < 0.1, green points) in SCZD vs. Control cases. Horizontal dashed line demarcates the y-axis value corresponding to the significance threshold FDR = 0.1. Vertical dashed lines demarcate log2 fold change values of -1 and 1. **E)** SCZD vs. Control DEG comparison between Hb (+Thal.) and four other brain regions: Caudate, DG, HIPPO, and DLPFC. Only genes expressed in the Hb were considered in the comparative analysis. X and Y axes are *t*-statistic values. Red points are unique DEGs (FDR < 0.05) in Hb (+Thal.) compared to other brain regions tested. Blue points are DEGs (FDR < 0.05) found in comparison regions, but not Hb (+Thal.). Purple points are DEGs (FDR < 0.05) that overlap between Hb (+Thal.) and comparison regions.

Prior to performing differential gene expression (DGE) analysis, we assessed how the transcriptomic landscape of our Hb-enriched samples compared to that of other brain regions and subregions implicated in psychiatric disorders. We conducted principal component analysis (PCA) on bulk RNA-seq data (subsetted to neurotypical control samples) from eleven regions (n = 817 total samples), including the amygdala, hippocampus, caudate, and several cortical areas (**Fig. 4B-C**). Hb-enriched samples separated from other brain regions’ samples along the sixth principal component axis (PC6), which explained 2.64% of the variance in the combined dataset, highlighting that Hb-enriched thalamus has unique transcriptional features compared to other brain regions.

We then performed DGE analysis by diagnosis (SCZD vs. Control samples) to identify genes significantly upregulated or downregulated (FDR < 0.1) in Hb-enriched tissue (**Fig. 4D; Table S7**). We controlled for effects of covariates including age, estimated proportions of Hb and thalamic cells in the samples, bulk RNA-seq quality control metrics, and quality surrogate variables associated with RNA degradation *(51, 52)*, all of which explained a portion of the gene expression variance (**Supplementary Methods, Fig. S17**). We found 173 differentially expressed genes (DEGs, FDR < 0.1) between SCZD cases and controls, including several genes previously implicated in SCZD, such as *NOTCH4* (**Fig. 4D**) *(37, 53–56)*. To evaluate SCZD-associated gene expression differences unique to Hb-enriched tissue compared to other brain regions, we conducted a cross-region comparison of DEGs obtained from our Hb-enriched dataset (n=45 DEGs, FDR < 0.05) and those from previously published bulk RNA-seq studies of four other brain regions: dorsolateral prefrontal cortex (DLPFC), hippocampus, dentate gyrus, and caudate *(19–21)* (**Fig. 4E**). We found the highest number of overlapping DEGs (n=12, FDR < 0.05) with the caudate, with one or less overlapping DEGs in other brain regions tested. *PHOSPHO1*, an upregulated DEG in both Hb-enriched samples and caudate, showed enrichment in the human LHb.3 subpopulation (enrichment *t*-stat = 4.415). In summary, we identified 32 DEGs (FDR < 0.05) between Control and SCZD samples that were unique to the Hb-enriched dataset compared to other brain regions. These results support a critical role for the human Hb in neuropsychiatric disease, and provide potential Hb-specific molecular targets for functional follow up and therapeutic applications.

### Hb eQTL identification

By combining single nucleotide polymorphism (SNP) DNA genotype data with bulk RNA-seq gene expression data, we identified independent expression quantitative trait loci (eQTL) for 717 SNP-gene pairs involving 707 unique genes and 687 unique SNPs (**Supplementary Methods**, **Table S8**) *(57)*. Seven pairs involved SCZD DEGs in Hb-enriched tissue (FDR<0.1, **Fig. 5A**, **Fig. S18**, Fisher’s exact test p=0.3045 for a positive association, **Table S9A**) and 16 different pairs involved SCZD GWAS risk SNPs (p<5 * 10^-8^, **Fig. 5B**, **Fig. S19**, Fisher’s exact test p=4.571e-08 for a positive association, **Table S9B**) *(58)*. By evaluating the effect of the estimated Hb or thalamus neuronal cell fractions (**Fig. S20**), 7 SNP-gene pairs have increased eQTL signal when Hb fraction increases and thalamus fraction decreases (**Fig. 5C**). Most independent eQTLs identified in Hb-enriched tissue were either unique or more strongly associated in Hb than in DLPFC or hippocampus (**Fig. 5D-E**) *(21)*. Colocalization analysis *(59)* of eQTL and SCZD PGC3 GWAS data identified colocalization for 16 genes (PP4 > 0.8), with 123 out of 256 (47.3%) SNPs from the 95% credible sets being SCZD risks SNPs (**Table S10**). *PCCB* (**Table S10**) was previously identified as a colocalized gene with SCZD in a multi-ancestry meta-analysis from 2,119 brain donors from other brain regions *(60)*. In the developing brain, *ANKRD45*, *CCDC122*, *DNAH10OS*, *LRRC37A2*, and *PCCB* (**Table S10**) were recently colocalized with SCZD *(61)*. *ANKRD45*, *ATXN7*, *GABBR2*, and *PCCB* (**Table S10**) are among a set of 1,111 SCZD risk genes involving different methodologies *(62)*, including eQTL colocalization, with *ANKRD45* and *ATXN7* included in 321 high confidence SCZD risk genes from an adult brain meta-analysis from 1,866 donors *(62)*. Thus 9 SCZD colocalized genes in Hb (56.3%) have not been previously identified by colocalization analyses in other brain regions, including *C5orf63*, *CD40*, *IFT52*, *PABPC1L*, *RGS16*, *RP11-166B2.1*, *SLC25A27*, *TDRD6*, and *UPF1* (**Table S10**). These results reveal eQTLs that are unique or enriched in Hb and provide evidence that the Hb contributes to the causative genetic architecture of SCZD.

**Fig. 5.**
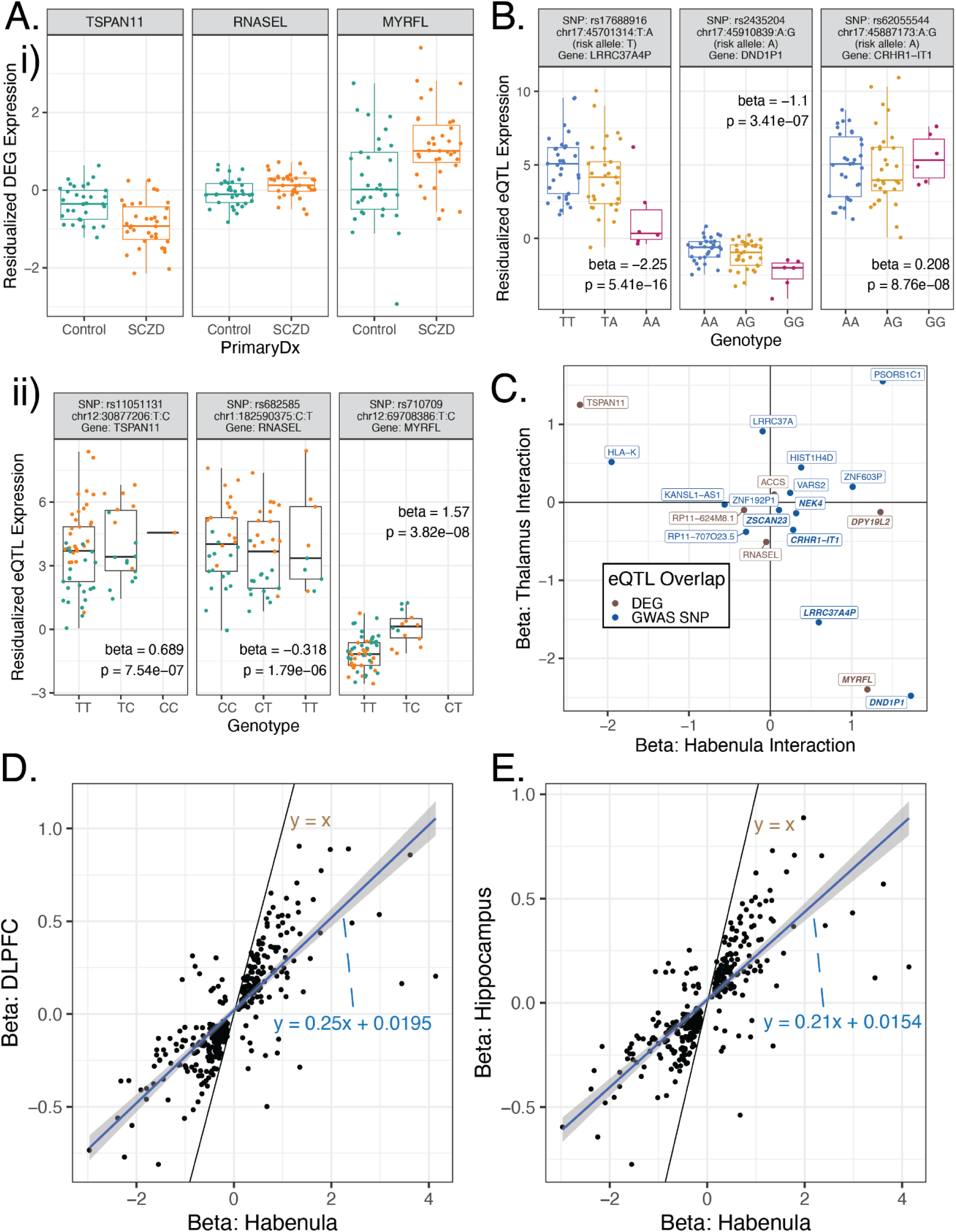
Identification of Hb-enriched thalamus independent expression Quantitative Trait Loci (eQTLs). **A)** Residualized normalized expression for SCZD DEGs (FDR<0.1) that are part of SNP-gene pairs identified as independent eQTLs (FDR<0.05). Boxplots by **i)** SCZD case-control status, or **ii)** SNP alleles. Aii) includes the eQTL regression (beta) and nominal association p-value. **B)** Boxplots by SNP alleles of residualized normalized expression for three genes from independent eQTLs (FDR<0.05) whose SNPs are PGC3 SCZD GWAS risk SNPs (p<5 * 10^-8^). Related to **Fig. S19**. Plots include eQTL regression (beta) and nominal association p-value. **C)** eQTL beta coefficients for the interaction between the SNP and the estimated habenula (`Tot.Hb`) or thalamus (`Tot.Thal`) neuronal cell proportions. All SNP-gene pairs either contain a SCZD DEG (FDR<0.1) or a PGC3 SCZD GWAS risk SNP (p<5 * 10^-8^), and are labeled with the paired genes’ symbols. Higher values indicate that the eQTL signal increases as the corresponding region neuronal cell proportion increases. Related to **Fig. S20**. **D)** eQTL beta coefficients for independent eQTLs in Hb-enriched thalamus compared against eQTL beta coefficients in BrainSEQ Phase II DLPFC data. Linear trend is shown in blue and identity line in brown. Points in black have stronger eQTL signal (beta coefficients) in Hb than in DLPFC, whereas points in gray have stronger or opposite signal in DLPFC. **E)** Same as D) but for comparing Hb-enriched thalamus against BrainSEQ Phase II hippocampus data. Out of the 717 Hb-enriched thalamus independent eQTLs, 324 and 317 of them were present in DLPFC (D) and hippocampus (E), respectively.

## Discussion

SCZD is a heritable polygenic disorder characterized by positive and negative symptoms with heterogeneity in clinical presentation and treatment response *(63)*. Therefore, it is important to understand how molecular changes in multiple brain regions contribute to SCZD etiology. Here, we focused on the Hb given its emerging role in psychiatric disorders and functional influence on neurotransmitter systems impacted in SCZD. To provide a basis for interpreting molecular changes associated with SCZD, we generated a topographic molecular map characterizing MHb and LHb cell type diversity in the healthy human brain. For instance, we identified a spatially-organized cholinergic cell type (MHb.2), which was distinguished from other *CHRNB4*-expressing MHb subpopulations by enrichment of *CHAT* and *CHRNA6*. This cholinergic subpopulation, which is conserved across species, may represent a shared biological substrate for comorbid nicotine dependence and SCZD *(64–68)*. Beyond classifying cell types in the human Hb, we also leveraged our reference atlas to disentangle molecular signatures from Hb and thalamus in a bulk RNA-sequencing study comparing SCZD and control donors.

Differential gene expression analysis of bulk RNA-seq data from individuals with SCZD and controls identified 32 differentially expressed genes (DEGs) unique to Hb-enriched tissue compared to other brain regions. Among unique DEGs, we identified downregulation of *HES5* (FDR < 0.05), an effector of Notch signaling highly expressed in neural stem cells (NSC) and implicated in the regulation of early-stage neurodevelopment. While Notch signaling is important for NSC fate, it is also implicated in neuroglial development *(53)*. Interestingly, we found *HES5* to be enriched in astrocytes (**Fig. 1**; enrichment t-stat = 7.794), suggesting this glial population in epithalamus may be particularly affected by downregulation of *HES5* in SCZD. In a mouse model of SCZD, *HES5* deficiency in cultured adult hippocampal cells affected the proliferative capacity of uncommitted progenitors, suggesting that *HES5* may also have important developmental functions in neurons *(69)*. Unique to Hb-enriched tissue, we also observed downregulation of dopamine receptor *DRD5* (FDR < 0.1). While little is known about the role of *DRD5* in SCZD, *DRD5* disruption in some animal models, but not others, is associated with cognitive deficits *(70–72)* and altered cortical oscillations *(73)*. Given the role of dopamine signaling in SCZD pathogenesis and treatment, future functional follow up studies should explore how *DRD5* signaling is altered in the Hb in the context of SCZD.

Cross-region comparison of SCZD-associated DEGs in Hb-enriched tissue with those identified in the caudate, DLPFC, HIPPO, and dentate gyrus (DG) revealed the greatest overlap with caudate DEGs. For instance, *NOTCH4* (FDR < 0.1) was upregulated in both Hb-enriched and caudate SCZD samples. *NOTCH4* is associated with genetic susceptibility to SCZD *(37, 54–56)*, and plays a key role in neural maturation and plasticity. Consistent with the neurodevelopmental hypothesis of SCZD etiology *(39)*, *Notch4* knockdown in a mouse NSC model leads to aberrations in cell proliferation, differentiation, and migration *(37–40)*. We also identified overlapping upregulation of *PHOSPHO1* (FDR < 0.05) in the Hb and caudate datasets. *PHOSPHO1* encodes a phosphatase that hydrolyzes phosphocholine to choline. Given that choline is necessary for synthesizing the neurotransmitter acetylcholine, this finding may suggest that cholinergic neurons and cell populations expressing cholinergic receptors, particularly MHb.2, are more vulnerable to SCZD pathology. Moreover, *PHOSPHO1* is regulated by a non-coding RNA (miR1306) in the chromosomal locus 22q11.2 *(74)*. Microdeletions in this locus cause DiGeorge syndrome and lead to cognitive and behavioral impairments, as well as a strikingly elevated risk of developing SCZD or schizoaffective disorder *(75–78)*. Further exploration of how PHOSPHO1 may be dysregulated in SCZD is warranted.

Independent eQTL results identified eQTL signals that are enriched in or unique to Hb. Hb-enriched independent eQTLs overlapped with SCZD DEGs and SCZD GWAS risk SNPs, highlighting the relevance of the Hb to genetic risk for SCZD. Of note is an independent eQTL (FDR < 0.05) overlapping with SCZD DEG *TSPAN11* (FDR < 0.1), as various genes of the tetraspanin membrane protein family have previously been implicated in neurodegenerative and neurodevelopmental disorders *(79–82)*, intellectual disability *(83–85)*, glutamatergic plasticity *(86–88)*, and regulation of inflammatory pathways *(89–91)*. Among SCZD GWAS risk SNPs overlapping with Hb-enriched independent eQTLs, the *LRRC37* gene family *(92, 93)* stands out due to its location on the chromosomal 17q21.31 locus. This locus contains other SNP-gene pairs we identified, including *DND1P1*, *CRHR1*, and *KANSL1* (**Fig. 5C**) and is associated with risk for neurodegenerative diseases, such as Parkinson’s disease (PD) and Alzheimer’s disease *(94–97)*, and neurodevelopmental disorders, such as autism spectrum disorder and SCZD *(98, 99)*. Of particular interest in this locus is *LRRC37A2,* a SCZD colocalized gene we identified in Hb which is also associated with PD by transcriptome wide analysis (TWAS) in multiple brain regions *(100)*. In addition to underlying a broad array of neuropsychiatric disease susceptibility *(101, 102)*, the 17q21.31 locus also has an association with cortical and subcortical morphology *(103–105)*. Finally, while eQTL and SCZD PGC3 GWAS colocalization results in Hb corroborate prior findings in developing and adult brains *(60–62)*, unique Hb results suggest that the Hb could play a pathogenic role in SCZD. Among overlapping results with previous studies *(60–62)*, *PCCB*’s contribution to SCZD risk has been recently studied in human forebrain organoids and its downregulation was linked to SCZD risk through the GABAergic pathways and mitochondrial function *(106)*. Future studies could investigate the function of *PCCB* in the etiology of SCZD in Hb.

Here we conduct the first transcriptomic study of postmortem human Hb in the context of SCZD. As with all postmortem human brain studies, we acknowledge our inability to confirm whether gene expression changes are the cause or consequence of disease. We tried to overcome some limitations by accounting for potential confounds related to donor demographics and RNA quality in our statistical model. We were also able to overcome challenges in precisely dissecting Hb tissue by performing cell type deconvolution of bulk RNA-seq data to control for thalamic contamination. Finally, we acknowledge that our study is limited to a small number of donors, including only Caucasian males, and that it will be important to replicate these findings in additional cohorts. In summary, we generated the first single cell transcriptomic atlas of the human Hb and used this reference dataset in conjunction with bulk RNA-sequencing and DNA genotyping of individuals with SCZD and neurotypical controls to better understand molecular changes associated with SCZD risk and illness state.

### Materials and Methods summary

Detailed materials and methods can be found in the supplementary materials. <u>Postmortem human brain tissue</u> Postmortem human brain tissue was collected from the Lieber Institute for Brain Development (LIBD) Human Brain and Tissue Repository through the following locations and protocols at the time of autopsy. Thirty-one samples were collected at LIBD between May 2013 and December 2017, twenty-eight of which were consented through the Office of the Chief Medical Examiner of the State of Maryland, under the Maryland Department of Health’s IRB protocol #12–24. The remaining three samples were consented through the Department of Pathology at Western Michigan University Homer Stryker MD School of Medicine, under WCG IRB protocol #20111080. Thirty-seven additional samples were consented through the National Institute of Mental Health Intramural Research Program (NIH protocol #90-M-0142), and were acquired by LIBD via material transfer agreement. Demographics and other information for the 69 donors are listed in **Table S1**. Details of tissue acquisition, handling, processing, dissection, clinical characterization, diagnoses, neuropathological examinations, and quality control measures are previously described *(107)*. Fresh frozen Hb-enriched epithalamic tissue samples were microdissected, pulverized, and stored at -80°C prior to snRNA-seq and bulk RNA-seq experiments. For single molecule fluorescent *in situ* hybridization (smFISH) experiments, larger tissue blocks were removed and kept intact for cryosectioning.

### Single nucleus RNA-sequencing (snRNA-seq) and analysis

Nuclei isolation was performed on seven neurotypical control tissue samples also included in the bulk RNA-seq study (**Table S1**) as previously described *(108)*. Nuclei were stained with propidium iodide (Invitrogen, Thermo Fisher) and sorted using fluorescence-activated nuclei sorting (FANS). Four samples were also stained with anti-NeuN (Millipore Sigma) for enrichment of neuronal nuclei. Using a BioRad S3e cell sorter, ∼9K nuclei were sorted per sample and loaded onto individual lanes of the 10x Genomics Chromium instrument. GEMs were produced, cDNA was generated, and libraries were prepared according to manufacturer’s instructions and sequenced on an Illumina Nextseq (**Table S2**). Reads were aligned with *CellRanger* (10x Genomics), empty droplets were identified with *DropletUtils (109)*, and nuclei quality was assessed by examining library size, number of detected genes, doublet scores, and fraction of reads mapping to mitochondrial and ribosomal genes (**Fig. S1-2**). Batch correction was performed with *Harmony (110)*, and dimension reduction with a generalized linear model for principal component analysis *(111)* (**Fig. S3-5**). Clustering was performed with graph-based clustering from *scran (112)*. Fine-resolution Hb clusters were maintained, but other captured cell types were collapsed into broader categories yielding 17 annotated cell type clusters containing 16,437 nuclei (**Fig. 1B**). Marker genes for each Hb subcluster were selected with the *Mean Ratio* method (**Fig. 1E**; **Table S3**).

### Single molecule fluorescent *in situ* hybridization (smFISH)

smFISH experiments were performed on postmortem human brain tissue using RNAScope Multiplex Fluorescent Reagent Kit v2 (Advanced Cell Diagnostics) and 4-Plex Ancillary Kit as previously described *(113)*. Briefly, Hb tissue from three independent donors (**Table S4**) was cryosectioned for four experiments targeting different Hb cell type clusters identified using our snRNA-seq dataset (**Fig. 1**, **Fig. S6A**, **Fig. S7A**, **Fig. S8A**, **Table S5**). Two probe combinations targeted LHb subclusters and two probe combinations targeted MHb subclusters (see **Supplemental Methods**). Briefly, tissue sections (n=2-4 technical replicates per donor) were fixed in 10% Normal Buffered Formalin (NBF) solution, rinsed in phosphate buffered saline (PBS), and dehydrated in a series of ethanol dilutions. Sections were treated with hydrogen peroxide, rinsed in PBS, and permeabilized with Protease IV. Following PBS wash steps, sections were hybridized with RNAScope probes, and signal amplification was performed according to manufacturer’s instructions. A distinct Opal fluorophore (Perkin Elmer, 1:500) was assigned to each probe and sections were counterstained with DAPI (4′,6-diamidino-2-phenylindole) to label nuclei. For each tissue section, a 20X max-intensity projected z-stack image was obtained with a Nikon AXR confocal microscope using spectral imaging and linear unmixing as previously described *(113)*. Confocal imaging data were saved as .nd2 files for downstream quantitative analysis using HALO software (Indica labs). The distributions of estimated marker gene expression (“copies”) across cell objects were used to quantify abundant expression (top ranks) and delineate thresholds for data display (**Fig. S9**).

### Bulk RNA-sequencing

Total RNA was extracted from tissue samples using the Qiagen AllPrep DNA/RNA/miRNA Universal Kit (Cat No./ID: 80224). Paired-end strand-specific sequencing libraries were prepared for 34 Control and 35 SCZD samples from 300 ng total RNA using the TruSeq Stranded Total RNA Library Preparation kit with Ribo-Zero Gold ribosomal RNA depletion. For quality control, synthetic External RNA Controls Consortium (ERCC) RNA Mix 1 (Thermo Fisher Scientific) was spiked into each sample. Libraries were sequenced on an Illumina HiSeq 3000 producing 30.14 to 645.8 million (median 90.65, mean 150.07) 100-bp paired-end reads per sample. Bulk RNA-seq FASTQ files were aligned to Gencode v25 *(114)* using *SPEAQeasy (115)*, a *Nextflow (116)* workflow for *HISAT2* RNA-seq alignments *(117)*. All samples passed quality control based on sequencing metrics (**Fig. S10**).

### Deconvolution

Given the small size of the Hb and inclusion of surrounding thalamic tissue during microdissection, cell type deconvolution of the bulk RNA-seq data was performed with *Bisque (118)*. Medial and lateral Hb cell type subclusters were collapsed and the top 25 mean ratio marker genes for each broad cell type were selected for deconvolution (**Table S6**). For validation purposes, we also calculated the standard log2 Fold Change for selected marker genes *(112)* with a model adjusting for the donor ID and contrasting a given cell type against all the rest (**Fig. S11**). One bulk RNA-seq sample (Br5572) was excluded from further analyses as it was predicted to consist entirely of thalamic tissue (**Fig. S12**).

### Cross-brain region bulk RNA-seq data integration

To examine the unique gene expression profile of the Hb, the control bulk RNA-seq samples from this study were compared to bulk RNA-seq samples from 11 other brain regions: Amygdala *(119)*, basolateral amygdala (BLA) *(120)*, central amygdala (CeA) *(121)*, caudate *(20)*, dorsal anterior cingulate cortex (dACC) *(120)*, dentate gyrus (DG) *(19)*, dorsolateral prefrontal cortex (DLPFC) *(21, 120)*, hippocampus (HIPPO) *(21, 120)*, medial amygdala (MeA) *(120)*, medial prefrontal cortex (mPFC) *(121)*, and subgenual anterior cingulate cortex (sACC) *(119)*. These LIBD samples came from donors with the same demographics: male, neurotypical, adult at age of death (17-70 years), and of European descent (EUR/CAUC). Principal components (PCs) were computed with log normalized gene expression (**Fig. 4B-C**).

### Bulk RNA-seq Differential Gene Expression (DGE)

Gene level differential expression analysis between SCZD and Control Hb-enriched bulk RNA-seq samples was performed using the limma-voom method *(122)* (**Fig. 4D**) while adjusting for demographics, quality control metrics, quality surrogate variables, and estimated proportions of Hb and Thalamus cell types in the samples *(51, 52)*. To assess the variability in SCZD-associated DEGs across brain regions, the Hb (+ Thal.) DGE results were compared to statistics from previous SCZD bulk RNA-seq studies: DLPFC, HIPPO *(21)*, DG *(19)*, and caudate *(20)* (**Table S7**, **Fig. 4E**). Bulk and snRNA-seq gene expression data from this study can be interactively explored through *iSEE*-powered websites https://github.com/LieberInstitute/Habenula_Pilot#interactive-websites *(123)*.

### Expression quantitative trait loci (eQTL) analysis

Gene level eQTL analysis was performed using a set of 5,504,021 SNPs with a minor allele frequency greater than 5% on a ±500kb window around genes with *tensorQTL (57)*. Nominal and independent eQTLs (FDR < 0.05) were computed on the log normalized gene expression values while adjusting for SCZD status, the top 5 ancestry PCs, and top 13 gene expression PCs (**Fig. 5**, **Table S8**). Interaction eQTLs (FDR < 0.05) with the neuronal cell proportions from Hb or Thal. were also identified.

### Colocalization analysis

Colocalization analysis was performed with *coloc (59)* with a posterior probability for H_4_ (PP4) greater than 0.8 on a ±500kb window around genes. Colocalized SNPs were then identified using a 95% credible set for H_4_.

## Acknowledgements

We gratefully thank the families who donated tissue to make this research possible. We thank the families of Connie and Stephen Lieber and Milton and Tamar Maltz for their generous support of this work. We thank Amy Deep-Soboslay and James Tooke for their work in sample curation and clinical characterization at LIBD. We thank Linda Orzolek and the Johns Hopkins Single Cell and Transcriptomics Core facility for executing the snRNA-seq sequencing. We thank the Joint High Performance Computing Exchange (JHPCE) for providing computing resources for these analyses. We thank Keri Martinowich (LIBD) for constructive feedback on the manuscript.

## Funding

This work was supported with funding from the Lieber Institute for Brain Development and National Institutes of Health grants T32MH015330 (Yalcinbas) and R01DA055823 (Maynard). Bulk RNA-seq sample selection and data generation was generated with funding from F. Hoffmann-La Roche AG.

## Author Contributions

Conceptualization: EAY, BA, DRW, LAHM, LCT, KRM

Methodology: LAHM, DRW, LCT, KRM

Software: LAHM, LCT

Validation: EAY, KDM, JW, HRD, AC, SB

Formal Analysis: EAY, BA, EDN, RGF, NJE, JMS, AC, LAHM, LCT

Investigation: EAY, EDN, KDM, JW, HRD, AC, RB, SB, RJ, RT, KRM

Resources: JHS, JEK, TMH, DRW, LCT, KRM

Data Curation: EAY, BA, RGF, GP, LAHM, LCT

Writing-original draft: EAY, BA, RGF, NJE, GP, KDM, LAHM, LCT, KRM

Writing-review and editing: EAY, LAHM, LCT, KRM

Visualization: EAY, BA, EDN, RGF, NJE, LAHM, LCT

Supervision: LAHM, DRW, LCT, KRM

Project administration: DRW, LCT, KRM

Funding Acquisition: DRW, LCT, KRM LCT and KRM had full access to all the data in the study and take responsibility for the integrity of the data and the accuracy of the data analysis.

## Competing Interests

Joel E. Kleinman is a consultant on a Data Monitoring Committee for an antipsychotic drug trial for Merck & Co., Inc.

## Data and materials availability

The source FASTQ files are publicly available from the Globus endpoints ‘jhpce#habenulaPilotbulkRNAseq’, ‘jhpce#habenulaPilotsnRNAseq’, and ‘jhpce#habenulaPilotRNAscope’ for the bulk RNA-seq, snRNA-seq, and RNAScope data, respectively. The DNA genotype data is available from ‘jhpce#habenulaPilotbulkDNAgenotype’ available upon request, given the protected nature of this data. All these Globus endpoints are listed at http://research.libd.org/globus. All source code developed for analyzing this data is available at https://github.com/LieberInstitute/Habenula_Pilot *(124)*.

## Materials and Methods

### Postmortem Human Clinical Characterization

Audiotaped and witnessed informed consent was obtained from the legal next-of-kin for every case. The LIBD Autopsy Phone Screening was performed at time of donation with the legal next-of-kin, and consisted of 39 items about the donor’s medical, social, psychiatric, substance use, and treatment history. Retrospective clinical diagnostic reviews were conducted for every brain donor, which included data from autopsy reports, toxicology testing, forensic investigations, neuropathological examinations, phone screening, and psychiatric/substance abuse treatment record reviews and/or supplemental family informant interviews. All data was compiled in a detailed psychiatric narrative summary, and was reviewed independently by 2 board-certified psychiatrists to determine lifetime psychiatric and substance use disorder diagnoses according to DSM-5. Every donor underwent toxicology testing by the medical examiner as part of the autopsy and forensic investigation for drugs of abuse such as ethanol/volatiles, cocaine/metabolites, amphetamines, and opiates. Additional supplemental toxicology testing was performed on all donors, including for nicotine/cotinine and cannabinoids. For psychiatric cases, supplemental testing for therapeutic drugs, such as antidepressants, mood stabilizing agents, and antipsychotics was completed through National Medical Services (www.nmslabs.com) in postmortem blood and/or cerebellar tissue. All non-psychiatric control donors had no lifetime history of a psychiatric or substance use disorder according to DSM-5.

### snRNA-seq Data Collection

Fresh frozen postmortem human brain tissue from seven adult neurotypical control male donors were dissected for epithalamus (Br1092, Br1204, Br1469, Br1735, Br5555, Br5558, and Br5639; see **Table S1** for demographics and other information). Tissue was homogenized and nuclei isolation was performed using the “Frankenstein” protocol as previously described *(108)*. All samples were stained with propidium iodide (Cat No. P3566, Invitrogen, Thermo Fisher, Waltham, MA). Additionally, four of the seven samples (Br1092, Br1204, Br5555, Br5558) were stained with Alexa Fluor 488-conjugated Anti-NeuN (Cat No. MAB377X, Millipore Sigma, St. Louis, MO) to enrich for neuronal nuclei during fluorescence-activated nuclei sorting (FANS). Using a BioRad S3e cell sorter, nuclei were sorted into 23.1 uL of master mix without enzyme C, prepared as per the 3’ next GEM Chromium Kit protocol (PN-1000075, 10x Genomics, Pleasanton, CA). Following FANS, GEMs were produced, cDNA was generated, and library preparations were completed by following revision A of the Chromium Next GEM Single Cell 3ʹ Reagent Kits v3.1 (Dual Index) protocol (PN-1000075, PN-1000073, CG000315, 10x Genomics). Samples were sequenced on an Illumina platform according to manufacturer’s instructions at the John Hopkins Single Cell and Transcriptomics Core.

### snRNA-seq Quality Control, Clustering, and Annotation

snRNA-seq samples were sequenced to a median depth of 193 million reads (min. 162.8, mean 188.3, max. 213), corresponding to a median 54,641 mean reads per nucleus (min. 39,476, mean 88,719, max. 230,795), a median of 14,135 median unique molecular indices (UMIs) per nucleus (min. 7,752, mean 17,674, max. 34,479), and a median 4,592 median genes per nucleus (min. 2,836, mean 5,083, max. 7,843). FASTQ files were aligned with *CellRanger* v6.0.0 (https://www.10xgenomics.com/support/software/cell-ranger/latest) against the `refdata-gex-GRCh38-2020-A` annotation distributed by 10x Genomics. The `--include-introns` option was used. Prior to quality control processing, there were 20,327 nuclei across all seven samples with a median of 3,156 nuclei per sample (min. 923, mean 2,904, max. 4,389) as estimated by *CellRanger* (**Table S2**).

Starting from the raw data instead of the *CellRanger* filtered data, empty droplets were excluded using *DropletUtils* v1.18.1 *(109)* with `emptyDrops(niters = 30000, lower = knee_lower)`, where `knee_lower` was determined for each sample by the “knee point” calculated by *DropletUtils* `barcodeRanks()` function plus 100. This resulted in `knee_lower` values ranging from 219 to 481. Droplets with a significant deviation from each sample’s ambient profile (FDR < 0.001) were kept, resulting in a dataset of 19,802 nuclei at this processing stage (936 to 3,905 per sample).

As part of quality control, nuclei were assessed for high mitochondrial content, low library size, and a low number of detected features. We applied an adapted 3 median absolute deviation (MAD) threshold per sample using *scater* v1.26.1 isOutlier(nmads=3)*(111)* (**Fig. S1**). Any nuclei that did not pass each threshold were excluded from further analyses (2,720 nuclei), resulting in 17,082 nuclei that passed this stage of quality control. Lastly, doublet scores were computed per sample using the top 1,000 highly variable genes with *scDblFinder* v1.12.0 `computerDoubletDensity()` *(125)*. This metric was later leveraged to assess the integrity of each cell type cluster. No cell type clusters were dropped as a result of their doublet scores.

Dimension reduction was performed using the generalized linear model for principal component analysis (GLM-PCA), using both *scry* v1.10.0 `nullResiduals()` *(126)* and *scater* v1.26.1 runPCA() *(111)*. It was apparent that the initial reduced dimensions showed evidence for batching by `Sample`, sequencing run, and NeuN sorting **(Fig. S3A, Fig. S4A, Fig. S5A)**. To correct for these batch effects, we applied *Harmony* v0.1.1 `RunHarmony(group.by.vars = “Sample”)` *(110)* to the GLM approximated PCs. This returned Harmony-corrected PCs, which showed evidence of successfully correcting for differences across the aforementioned variables **(Fig. S3B, Fig. S4B, Fig. S5B)**.

After correcting by `Sample`, we applied graph-based clustering measures leveraging *scran* v1.26.2 *(112)* `buildSNNGraph(k = 10)` and *igraph* v1.4.2 `cluster_walktrap()` *(127)*, generating 37 fine-resolution nuclei clusters. Using established marker genes *(23)*, we annotated our 37 fine-resolution nuclei clusters for cell type identities. This process yielded 17 cell type categories that maintained the distinct Hb subclusters while collapsing all other non-Hb clusters into their respective cell type populations (**Fig. 1B**): 3 Astrocyte clusters, 11 Excitatory Thalamus clusters, 5 Inhibitory Thalamus clusters, and 3 Oligodendrocyte clusters were collapsed. We were thus ultimately left with 7 broad cell type clusters (i.e. Oligodendrocytes, OPC, Microglia, Astrocytes, Endothelial, Inhibitory Thalamus, and Excitatory Thalamus) alongside our 7 Lateral Habenula (LHb) and 3 Medial Habenula (MHb) cell type subclusters.

No given cell type cluster was driven by doublet scoring as no cluster had a particularly high median doublet score (**Fig. S2A**). Out of the 17,082 nuclei post-`isOutlier()` filters, one small cluster (51 nuclei) did not have a clear cell type identity, but did have high expression of *SNAP25*. This small cluster was thus classified as “Excit.Neuron” and excluded from downstream analyses (**Fig. S2**). Nuclei from three donors (Br5555, Br1204, Br1092) in the OPC cluster were labeled as “OPC_noisy” (594 nuclei) given their less defined spatial arrangement in *t-*SNE dimensions 1 and 2, and were excluded from further analyses. The entire quality control, clustering, and annotation processes brought the initial number of 20,327 nuclei to a final total of 16,437 filtered and annotated nuclei (**Fig. S13**).

### snRNA-seq Marker Gene Selection

Marker genes were selected for the 17 identified cell type categories through the *Mean Ratio* method from *DeconvoBuddies*v0.99.0 (https://github.com/LieberInstitute/DeconvoBuddies), using the function `getMeanRatio2()`. The mean ratio method calculates, for each gene, the mean expression of a target cell type divided by the highest mean expression of a non-target cell type. A high mean ratio value for a gene in a given cell type cluster suggests that that gene is a cell type specific marker gene candidate for that cell type cluster. The top 50 mean ratio marker genes for each of our cell type categories are listed in a table (**Table S3**).

### Cross-species Comparison

Cell type clusters observed in the human snRNA-seq dataset were compared to a previously annotated single cell RNA-sequencing dataset from mouse habenula *(22)*. The mouse dataset had annotations across all cell types, and lateral and medial specific annotations for just the Hb neurons. To relate the mouse and human datasets, homologous gene IDs were found using the Mouse Gene Informatics Website. Both of the datasets were subset to 14,468 genes with valid homologs *(128)*. The two datasets were compared via spatial registration pipeline from *spatialLIBD* v1.12 *(129)*, and gene enrichment statistics for each cell type in both annotations were calculated with `registration_wrapper()`. For a more targeted comparative analysis, we utilized a “neuron-only” mouse dataset, and in this case, enrichment statistics were computed on the “habenula neuron-only” subset of our human snRNA-seq dataset. In both iterations of the comparative analyses, correlation values between the *t*-statistics were calculated with `layer_stat_cor(top_n = 100)` and heatmaps were plotted with *ComplexHeatmap* v2.16 *(130)* (**Fig. 2A-B**).

### Multiplexed Single molecule fluorescent *in situ* Hybridization (smFISH)

smFISH experiments were performed according to manufacturer’s instructions as previously described using the RNAScope Multiplex Fluorescent Reagent Kit v2 (Advanced Cell Diagnostics, Hayward, California, Cat No. 323100) and 4-Plex Ancillary Kit for Multiplex Fluorescent Kit v2 (Cat No. 323120) *(113)*. Fresh frozen tissue blocks containing Hb from three independent donors (**Table S4**) were cryosectioned at ∼10 μm on a Leica cryostat and stored at -80°C. Each donor yielded 2-3 tissue slides with 2-4 tissue sections per slide. Slides were assigned to three different RNAScope experiments targeting different LHb and MHb cell type clusters based on results from snRNA-seq (**Fig. 1**, **Fig. S6A**, **Fig. S7A**, **Fig. S8A**). <u>Experiment 1</u> <u>targeting LHb subclusters:</u> *ONECUT2* (Cat No. 473531-C1), *TLE2* (Custom Design 1271611-C2), *SEMA3D* (Cat No. 521771-C3), and *POU4F1* (Cat No. 438441-C4). <u>Experiment 2</u> <u>LHb subclusters:</u> *ESRP1* (Cat No. 435051-C1), *MCOLN3* (Cat No. 516761-C2), *CRH* (Cat No. 473661-C3), and *POU4F41* (Cat No. 438441-C4). <u>Experiment 3 MHb subclusters:</u> *CCK* (Cat No. 539041-C1)*, POU4F1* (Cat No. 438441-C2), *EBF3* (Cat No. 581641-C3), and *CHAT* (Cat No. 450671-C4); <u>or</u> *CCK* (Cat No. 539041-C1)*, CHRNB4* (Cat No. 482411-C2), *BHLHE22* (Cat No. 448351-C3), and *CHAT* (Cat No. 450671-C4) Briefly, tissue sections were fixed in 10% Normal Buffered Formalin (NBF) solution for 30 minutes at room temperature. Sections were then rinsed in 1x phosphate buffered saline (PBS) and sequentially dehydrated for 5 minutes each in four ethanol dilutions: 50%, 75%, 100%, and 100%. Once the tissue sections were dry, a hydrophobic pen was used to carefully outline them on the slides to form a hydrophobic barrier. Tissue sections were then treated with hydrogen peroxide for 10 minutes at room temperature. Following subsequent decanting and rinsing in 1xPBS, tissue sections were permeabilized with Protease IV for 30 minutes at room temperature. After decanting and rinsing the tissue slides in 1xPBS, probe hybridization solution was prepared. Three different sets of RNAScope probe combinations (4 probes per combination) were used for each experiment as described above.

Tissue sections were incubated in probe hybridization solution at 40°C for 2 hours. Following decanting and wash steps in 1x wash buffer, sections were incubated in saline-sodium citrate buffer (SSC) overnight at 4°C. Next, probe signal amplification steps were performed and a distinct Opal fluorophore (520, 570, 620, 690 nm) was assigned to each probe (Perkin Elmer, Waltham, MA; 1:500) (**Table S5**). Sections were counterstained with DAPI (4′,6-diamidino-2-phenylindole) to label nuclei. For each donor and probe combination, a 20X max-intensity projected z-stack image of the tissue section containing the largest habenula region (as gauged by signal in the *POU4F1* probe channel) was obtained with a Nikon AXR confocal microscope using spectral imaging and linear unmixing as previously described *(113)*. Confocal imaging data were saved as .nd2 files for downstream quantitative analysis.

### smFISH Confocal Image Analysis using HALO

HALO (Indica Labs) was used to segment and quantify fluorescent signals for each probe in single cells. Nikon .nd2 files were imported into HALO and an analysis magnification value of 2 (corresponding to 40X for a 20X image) was chosen as this is the recommended magnification for punctate probe signal quantification. For each image, nuclear detection and smFISH probe detection parameters were optimized to minimize false positives by referencing the *HALO 3.6 User Guide* (Indica labs, February 2023). For example, contrast threshold, minimum signal intensity, size (min and max), roundness (min), and segmentation aggressiveness were determined for each probe as well as DAPI. All HALO settings files are available on Github at https://github.com/LieberInstitute/Habenula_Pilot/tree/master/processed-data/14_RNAscope/HA <u>LO_data</u> *(124)*.

The *FISH-IF* module was used to quantify RNA transcripts (copy counts) within each detected object (i.e. a nucleus with dilated boundary to estimate a “cell”) with reference to the manufacturer’s guidelines: *HALO 3.3 FISH-IF Step-by-Step guide* (Indica labs, v2.1.4 July 2021). Distributions of object signal intensity values across the 4 RNAScope probe channels were assessed to determine representative values for each channel’s copy intensity parameter. The copy intensity parameter determines how many RNA transcript copies are assigned to each cell. For each of the RNAScope probe channels, the median of the probe signal intensity values of cells that had non-zero signal in that particular channel was chosen as a representative/typical intensity value for that channel’s copy intensity parameter. Thorough comparisons between the raw image files and HALO analysis segmentation outputs were conducted to ensure accurate representation of the imaging data.

### Visualization of Quantified smFISH Data

The spatial expression patterns of chosen probes as quantified by HALO were observed using hexbin plots displaying the maximum number of transcript copies in a bin generated with `stat_summary_hex(x = XMax, y = YMax, z = copies), fun = max, bins = 100)` from *ggplot2* v3.4 *(131)* (**Fig. S6C**, **Fig. S7C**, **Fig. S8C**). To focus on cell objects with robust expression of the marker gene probes, we visualized the top 100 cells ranked by number of transcript copies for each marker gene as a proxy for cell type. Note that ranks could include ties, in some cases leading to more than 100 cells being selected. As shown via confusion matrices, there was ∼10% overlap between most of the top 100 assignments depending on the donor slide and marker gene pair being compared (**Fig. S6B**, **Fig. S7B**, **Fig. S8B**). To observe the relative spatial location of the top 100 cells for each marker gene, points colored by the marker gene (and related Hb subcluster) identity were plotted in the X-Y locations of the cell objects, over a background of black points identifying high expression of the established Hb-wide marker gene *POU4F1*. If a cell had more than one top 100 identity, its point color reflected the marker gene with the maximum number of transcript copies in that cell (**Fig. 3A-Ci**, **Fig. S6B**, **Fig. S7B**, **Fig. S8B**).

### Bulk RNA-seq Data Collection

Total RNA was extracted from samples using the Qiagen AllPrep DNA/RNA/miRNA Universal Kit (Cat No./ID: 80224). Paired-end strand-specific sequencing libraries were prepared from 300 ng total RNA using the TruSeq Stranded Total RNA Library Preparation kit with Ribo-Zero Gold ribosomal RNA depletion (https://www.illumina.com/products/selection-tools/rrna-depletion-selection-guide.html) which removes rRNA and mtRNA. For quality control, synthetic External RNA Controls Consortium (ERCC) RNA Mix 1 (Thermo Fisher Scientific) was spiked into each sample. The libraries were sequenced on an Illumina HiSeq 3000 at the LIBD Sequencing Facility, producing from 30.14 to 645.8 million (median 90.65, mean 150.07) 100-bp paired-end reads per sample.

### Bulk RNA-seq Data Processing

Bulk RNA-seq FASTQs were aligned to Gencode v25 *(114)* using *SPEAQeasy*’s *(115)* development version that consisted of SGE scripts (https://github.com/LieberInstitute/RNAseq-pipeline). The settings were: `--experiment “Roche_Habenula” --prefix “PairedEnd” --reference “hg38” --stranded “reverse” --ercc “TRUE”`. This resulted in 24.3 to 612.18 million reads mapped (median 80.53, mean 132.48) with an overall mapping rate (overallMapRate) of 0.5471 to 0.9169 per sample (median 0.8506, mean 0.8368). All *SPEAQeasy* metrics are available (**Table S1**).

### Deconvolution

Cell type deconvolution of the bulk RNA-seq data was performed with Bisque *(118)* from the R package *BisqueRNA* version 1.0.5, with the function `ReferenceBasedDecomposition(use.overlap = FALSE)`. The snRNA-seq data from the present study (**Fig. 1**) was used as the reference dataset at the broad resolution; that is, by re-labeling MHb.1, MHb.2, MHb.3 as MHb, and similarly re-labeling the LHb fine-resolution clusters broadly as LHb. Across the broader-resolution clusters, marker genes were selected through the *Mean Ratio* method from *DeconvoBuddies* v0.99.0 (https://github.com/LieberInstitute/DeconvoBuddies) using the function `getMeanRatio2()`. The top 25 mean ratio marker genes for each cell type were selected as the marker genes for deconvolution (**Table S6**). For comparative purposes, we also calculated the standard log2 Fold Change (std.logFC, **Fig. S11**) using `findMarkers(test="t", direction="up", pval.type="all")` from *scran* v1.26.2 *(112)* with a model adjusting for the donor ID and contrasting a given cell type against all the rest. The top 25 mean ratio marker genes typically have a high standard log2 fold change (**Fig. S11**, **Table S6**) and can be visually inspected for validation (**Fig. S16**) using *iSEE (123)* interactive websites at https://github.com/LieberInstitute/Habenula_Pilot#interactive-websites.

### Bulk RNA-seq Quality Control

Sample Br5772 resulted in an estimated 100% inhibitory thalamus proportion, and was dropped from further analyses (**Fig. S12**). *SPEAQeasy (115)* quality control metrics such as the mitochondrial mapping rate (mitoRate) and ribosomal RNA mapping rate (rRNA rate) were visually inspected across each flow cell (**Fig. S10**), as well as by schizophrenia (SCZD) case-control status. Outlier samples in principal component analyses did not seem to be related to any specific metrics, except for Br5572, which as noted previously had unusual deconvolution results. Contrasting the range of these *SPEAQeasy* metrics against other similar studies *(19–21, 119)*, the remaining 68 bulk RNA-seq samples were considered of appropriate quality for downstream analyses.

### Bulk RNA-seq quality Surrogate Variables Calculation

To adjust for RNA degradation effects *(51)*, we calculated quality surrogate variables (qSVs) with *qsvaR (52)*. We used `qSVA(type = “standard”)`from *qsvaR* v1.5.3 on the filtered and normalized transcript data (TPM) with the model `∼ PrimaryDx + AgeDeath + Flowcell + mitoRate + rRNA_rate + totalAssignedGene + RIN + abs_ERCCsumLogErr + tot.Hb + tot.Thal`, where `tot.Hb = LHb + MHb` and `tot.Thal = Inhib.Thal + Excit.Thal`. This resulted in 8 qSVs from 1,772 transcripts associated with RNA degradation in six brain regions as described in the *qsvaR* documentation (**Table S1**). Note that habenula is not one of these six regions.

### Bulk RNA-seq Variance Partition Across Genes

We used both *scater* and *variancePartition* to explore the percent of variance explained across all genes (**Fig. S17**) by different *SPEAQeasy (115)* metrics, sample demographic variables such as age at time of death, and qSVs (**Table S1**). We ran `plotExplanatoryVariables()` from *scater* v1.28.0 *(111)* with default parameters. We also used `getVarianceExplained()`, `canCorPairs()`, and `fitExtractVarPartModel()` from *variancePartition* v1.30.2 *(132)* with the log normalized counts (`logcounts`). According to the *variancePartition* user guide, categorical variables such as `PrimaryDx` and `Flowcell` were treated as random effects.

### Bulk RNA-seq Differential Expression Analysis

Gene level differential expression analysis was performed using `calcNormFactors()` from *edgeR* v3.42.4 *(133)* and `voom()`, `lmFit()`, `eBayes()`, and `topTable()` from *limma* v3.56.2 *(122)* (**Table S7**). The design model was `∼ PrimaryDx + AgeDeath + Flowcell + mitoRate + rRNA_rate + totalAssignedGene + RIN + abs_ERCCsumLogErr + tot.Hb + tot.Thal + qSV[1-8]`. This model tested for differences by SCZD case-control status stored in the primary diagnosis (`PrimaryDx`) variable. Differential expression results were visualized using *EnhancedVolcano* v1.18.0 *(134)*.

### Cross-Brain Region bulk RNA-seq Data Integration

Control bulk RNA-seq Hb-enriched samples were compared to bulk RNA-seq samples generated at the Lieber Institute for Brain Development from other brain regions with the same demographics: neurotypical individuals, adults at age of death (17-70 years), male, and european descent (EUR/CAUC).

- Amygdala: n = 140, (Moods) *(119)*
- Basolateral amygdala (BLA): n=54, VA_PTSD *(120)*
- Central amygdala (CeA): n=11, PTSD BrainOmics *(121)*
- Caudate: n = 82, *(20)*
- Dorsal anterior cingulate cortex (dACC): n = 55, VA_PTSD *(120)*
- Dentate gyrus (DG): n = 41, DG Astellas *(19)*
- Dorsolateral prefrontal cortex (DLPFC): n = 121 (BSP2 + VA_PTSD) *(21, 120)*
- Hippocampus (HIPPO): n = 26 (BSP2 + VA_PTSD) *(21, 120)*
- Medial amygdala (MeA): n = 55, VA_PTSD *(120)*
- Medial prefrontal cortex (mPFC): n = 57, (BSP4+5 + PTSD BrainOmics) *(121)*
- Subgenual anterior cingulate cortex (sACC): n = 142 (Moods) *(119)*

Principal components were computed using `prcomp()` on log2(RPKM + 1) expression values for genes with a mean RPKM > 0.1 across all brain regions. The percent of variance explained was computed with `getPcaVars()` from *jaffelab* v0.99.31 (https://github.com/LieberInstitute/jaffelab).

### SCZD vs. Control Differential Gene Expression (DGE) Signal Comparison Across Brain Regions

We downloaded the schizophrenia vs. control DGE results from the BrainSEQ Phase II dorsolateral prefrontal cortex (DLPFC) and hippocampus (HIPPO) *(21)*, BrainSEQ Phase III caudate *(20)*, and dentate gyrus (DG) bulk RNA-seq datasets *(19)*. All of the above analysis results were generated at the Lieber Institute for Brain Development. We compared these results with the SCZD vs. Control DGE results from our Hb-enriched dataset by subsetting the BrainSEQ and DG datasets to only include genes that were expressed in Hb (**Table S7**, **Fig. 4E**).

Only 1 out of 236 DLPFC, 1 out of 46 HIPPO, 0 out of 10 DG, and 12 out of 2,437 caudate DEGs (FDR < 5%) were also significantly differentially expressed in Hb (FDR < 5%).

### Single Nucleotide Polymorphism (SNP) DNA Genotyping

Genotype calling was performed using several Illumina SNP BeadChip microarrays (4 samples with HumanMap650Y, 27 with Human1M-Duo, 7 with HumanOmni5-Quad, and 31 with Infinium Omni2.5-8). Using *Plink* v1.9 *(135)* we initially excluded variants with a minor allele frequency (MAF) less than 0.5%, a missingness rate of 5% or more, and a Hardy Weinberg equilibrium (HWE) p<1x10^-5^. Genotype phasing and imputation was performed on batches for each of the different SNP arrays from the LIBD brain bank using the TOPMED hg38 service and reference panel *(136)*, and each imputation batch was filtered to keep only variants with R-square (imputation quality information value) R2 > 0.9 and MAF > 0.05. We then extracted the 69 samples from these filtered imputation batches and merged them into a single genotype file, of 7,900,749 variants. This merged imputed genotypes file was finally filtered with *plink* to remove variants with missing call rates exceeding 5% (`--geno 0.05`), HWE below 1x10^-6^ (`--hwe 1e-6`) and MAF above 5% (`--maf 0.05`) for the remaining variants. This resulted in a final filtered file consisting of 5,504,021 common variants for these 69 donors.

### eQTL analysis

We used *tensorQTL* v1.0.8 *(57)* to identify SNP-gene pairs where the SNP was significantly associated with the gene expression using a linear regression model that adjusted for `PrimaryDx` and the first five SNP-based principal components. This analysis was performed for all 5,504,021 SNPs with a MAF>5% on a ±500kb window around genes on the 68 donors that passed RNA-seq QC. The nominal expression Quantitative Trait Loci (eQTLs) results were then filtered to first identify cis-eQTLs, and then independent eQTLs with `tensorqtl.cis.map_cis()` and `tensorqtl.cis.map_independent()` functions in *tensorQTL*, respectively (**Fig. 5**). The analysis was performed with normalized gene expression (`logcounts`) adjusting for the following covariates: `∼ PrimaryDx + snpPC[1-5] + genePC[1-13]`. `snpPC`s were computed on the DNA genotype data while `genePC`s were computed on the gene expression data.

Psychiatric Genomics Consortium (PGC) v3 data for the SCZD GWAS *(58)* was obtained from `PGC3_SCZ_wave3.european.autosome.public.v3.vcf.tsv.gz` *(137)* and filtered to SNPs with a p<5 * 10^-8^ SCZD risk association (**Fig. 5B**, **Fig. S19**). Risk alleles were determined using frequency of SCZD and NTC donors in the GWAS data. More specifically, if `BETA` > 0 (that is `FCAS` > `FCON`), the risk allele is `A1`, otherwise it is `A2`. Based on the documentation of the data, `A1` is the SNP reference allele, `A2` is the SNP alternate allele, `BETA` is the beta or ln(odds ratio) of A1 for SCZD risk, `FCAS` is the frequency of A1 in SCZD cases, and `FCON` is the frequency of A1 in controls. Note that 14,105 out of 20,446 (68.99%) SCZD risk SNPs (p<5 * 10^-8^) have risk alleles that are the reference alleles. Fifteen out of sixteen (93.75%) SNPs overlapping SCZD risk SNPs also have the risk allele matching the reference allele (**Fig. 5B**, **Fig. S19**).

Cell fraction interaction eQTLs were performed against the habenula (`Tot.Hb`) and thalamus (`Tot.Thal`) estimated neuronal fractions from the deconvolution analysis. Interaction eQTLs were identified by supplying `DataFrame`s of habenula and thalamus fractions to the `interaction_df` parameter in separate calls to the `tensorqtl.cis.map_nominal()` function in *tensorQTL* (**Fig. S20**). Slope of the interaction term were extracted from the `b_gi` output column as described by *tensorQTL*at https://github.com/broadinstitute/tensorqtl/blob/master/docs/outputs.md and compared between habenula and thalamus fractions (**Fig. 5C**). SNP-gene pairs with genes *BTN3A2, HLA-DMA* and *SMAGP* (**Fig. S20**) had no interaction eQTL results with *tensorQTL* (**Fig. 5C**).

BrainSEQ Phase II dorsolateral prefrontal cortex (DLPFC) and hippocampus (HIPPO) *(21)* eQTLs were used to compare the eQTL regression betas against Hb-enriched thalamus independent eQTL results (**Fig. 5D-E**).

### Colocalization analysis

Colocalization of eQTLs and PGC3 SCZD risk SNPs was performed with *coloc* v5.2.3 *(59)*. The set of nominal *tensorQTL* gene-SNP pairs *(57)*, which considered SNPs within a 500kb radius of each gene, were filtered to those whose SNPs observed in the PGC3 SCZD dataset *(58)*. Each such 1Mb region corresponded to a pair of datasets on which to invoke `coloc.abf()` with default arguments. For the eQTL-based dataset, `beta` and `varbeta` were defined by *tensorQTL* columns `slope` and `slope_se` squared; `type` was set to `“quant”` and `sdY` set to 1 (to reflect normalization internal to *tensorQTL* which standardizes the trait variance to 1). For the PGC3-based dataset, `beta` and `varbeta` were defined by columns `BETA` and `SE` squared; type was set to `“cc”` and `s` was defined as 0.409, the median value of `NCAS / (NCAS + NCON)` across all SNPs. We considered a gene to be colocalized if `PP.H4.abf`, from the output summary of `coloc.abf()`, exceeded 0.8. This is the estimated posterior probability for H_4_ (PP4): both traits are associated and share a single causal variant. A SNP was considered to be colocalized if it was in the 95% credible set for H_4_.

### Software

*ggplot2* v3.4.2 and earlier versions *(131)*, *R* versions 4.1, 4.2, and 4.3 *(138)*, and *Bioconductor* versions 3.14, 3.16, and 3.18 *(139)* were used for the analyses.

## Supplementary Figures

**Fig. S1.**
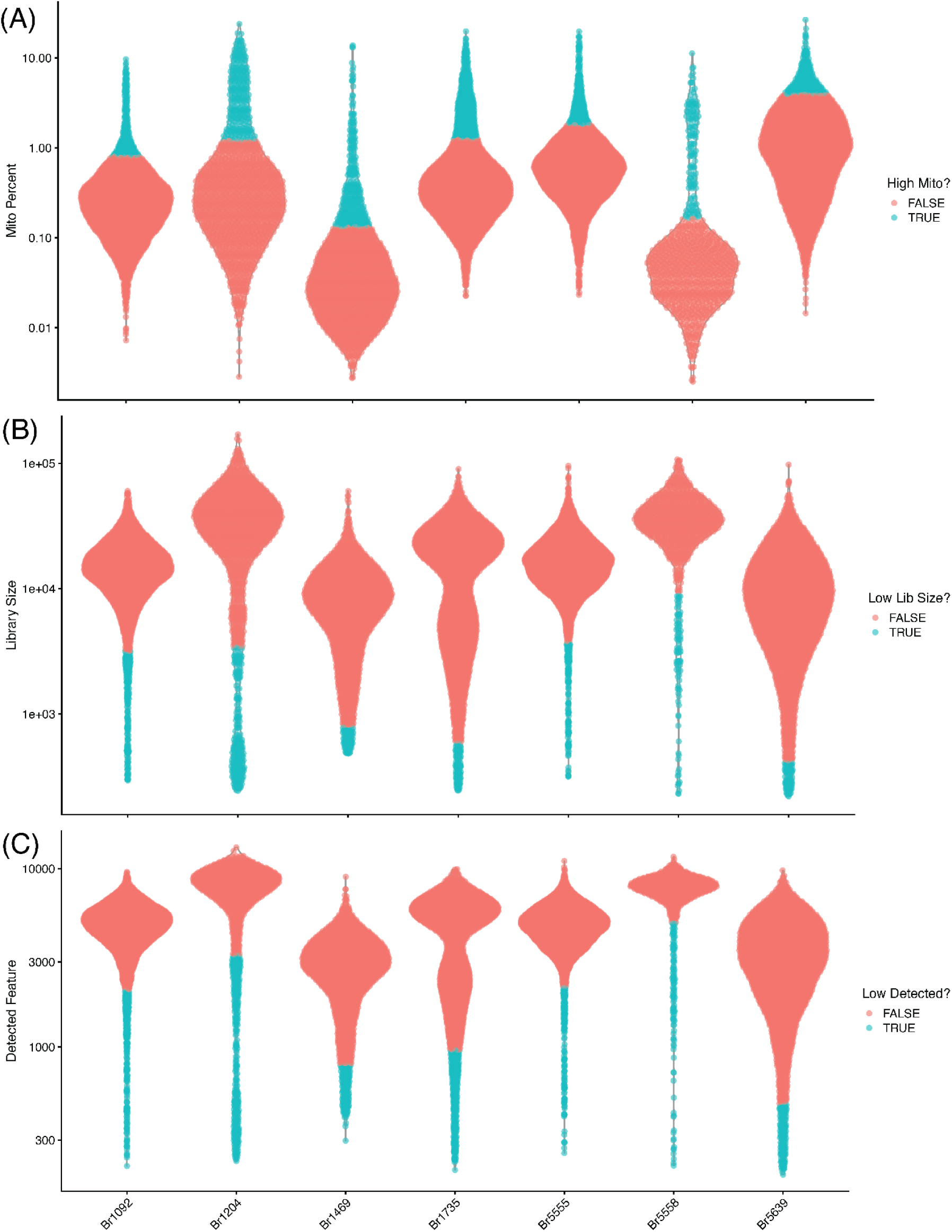
snRNA-seq quality control metrics. Violin distribution plots for **A**) mitochondrial percent, **B**) library size (total number of UMIs), and **C**) number of detected genes. Thresholds for quality control were determined using `isOutlier()` from *scuttle*.

**Fig. S2.**
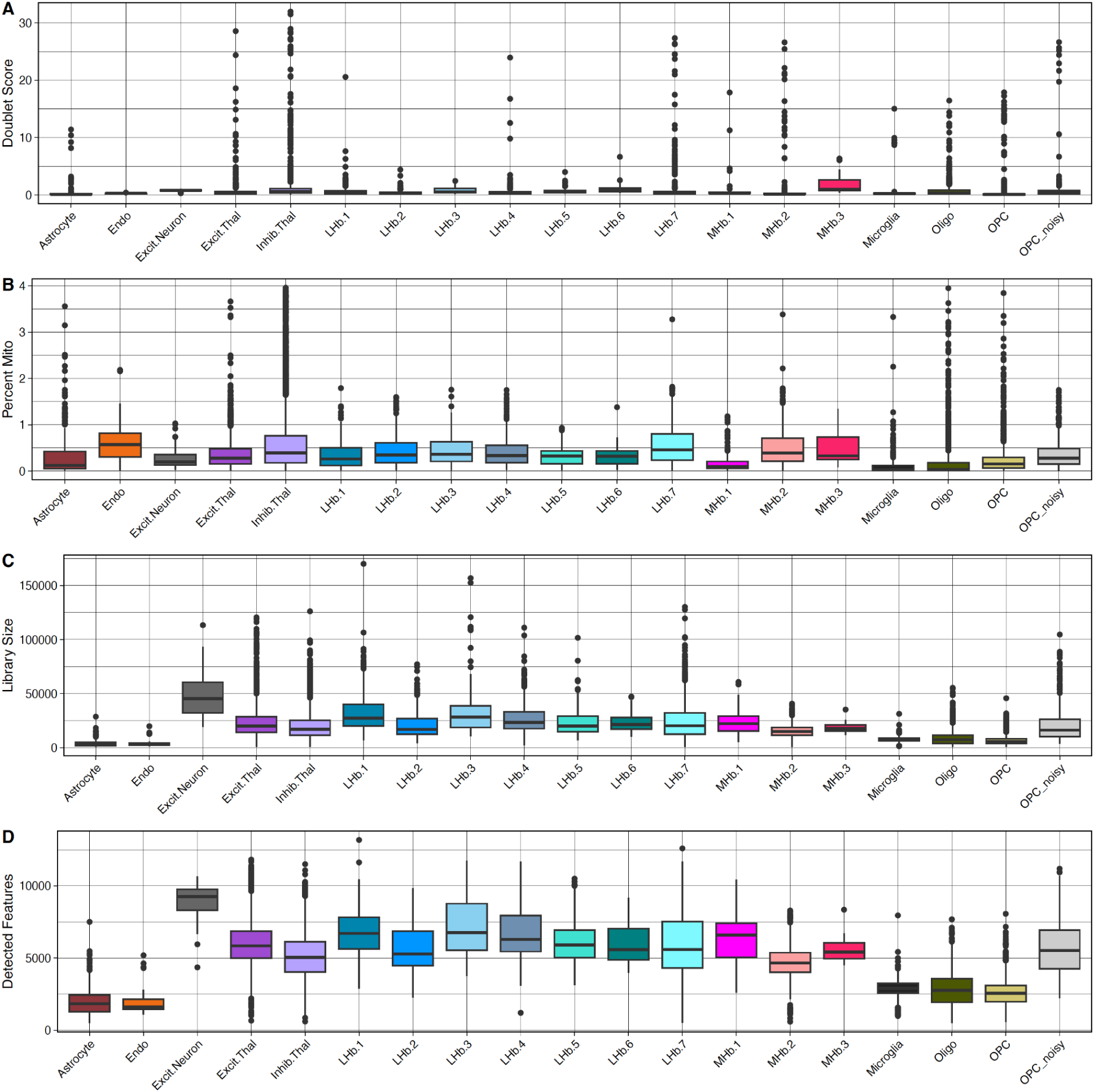
snRNA-seq quality control metrics by cell type. Boxplot distribution plots for each identified cell type looking at **A**) doublet score, **B**) mitochondrial percent, **C**) library size (total number of UMIs), and **D**) number of detected genes. Ambiguous excitatory neurons (Excit.Neuron) and OPCs (OPC_noisy) were dropped from further analyses.

**Fig. S3.**
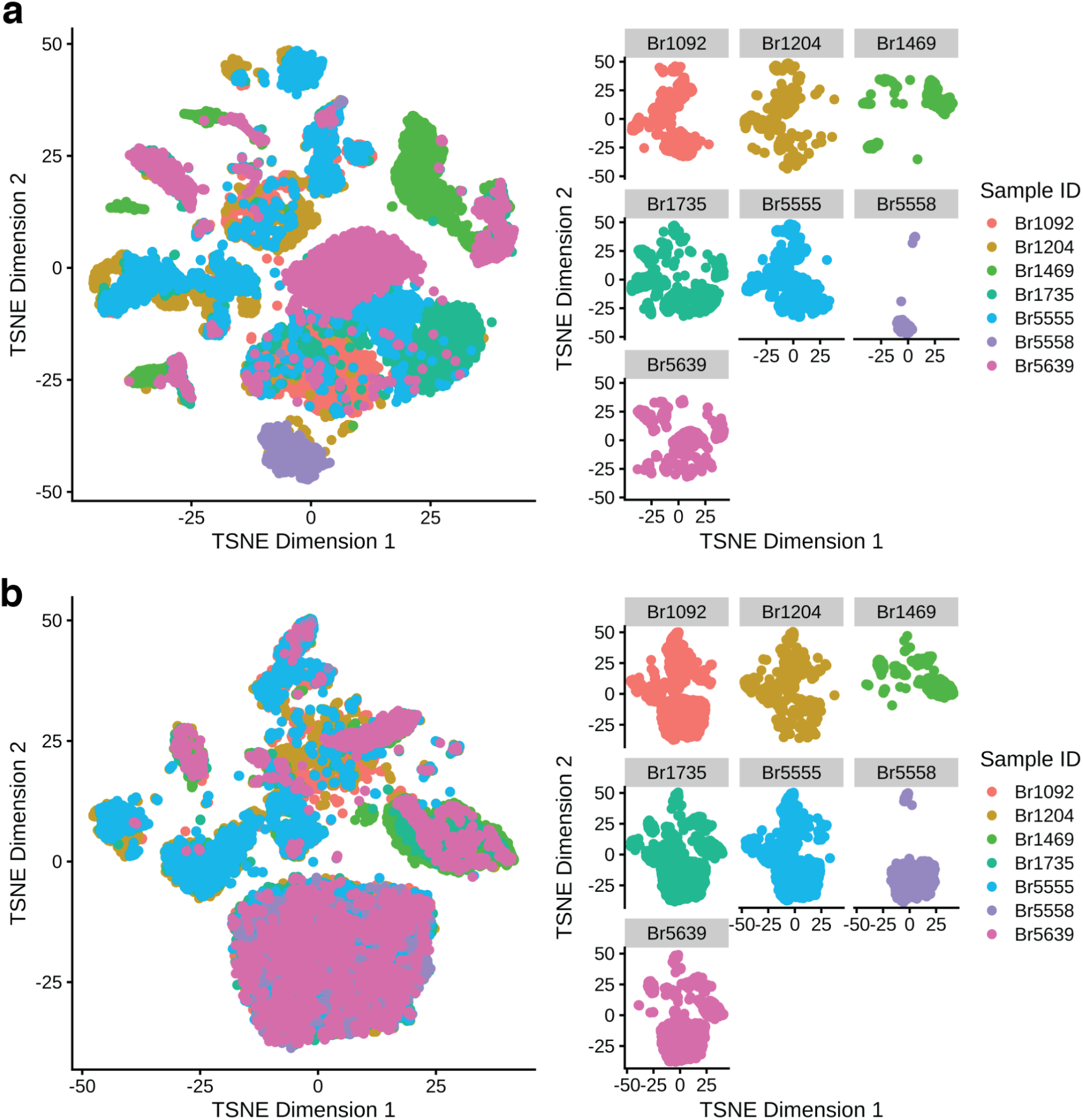
Pre and post harmony by sample. Batch correction of snRNA-seq data visualized by t-SNE plots. **A**) t-SNE of principal components (PCs) pre-batch correction, colored by sample. **B**) t-SNE of PCs post-batch correction with *Harmony*, colored by sample. Batch correction with *Harmony* reduced the sample batch effect.

**Fig. S4.**
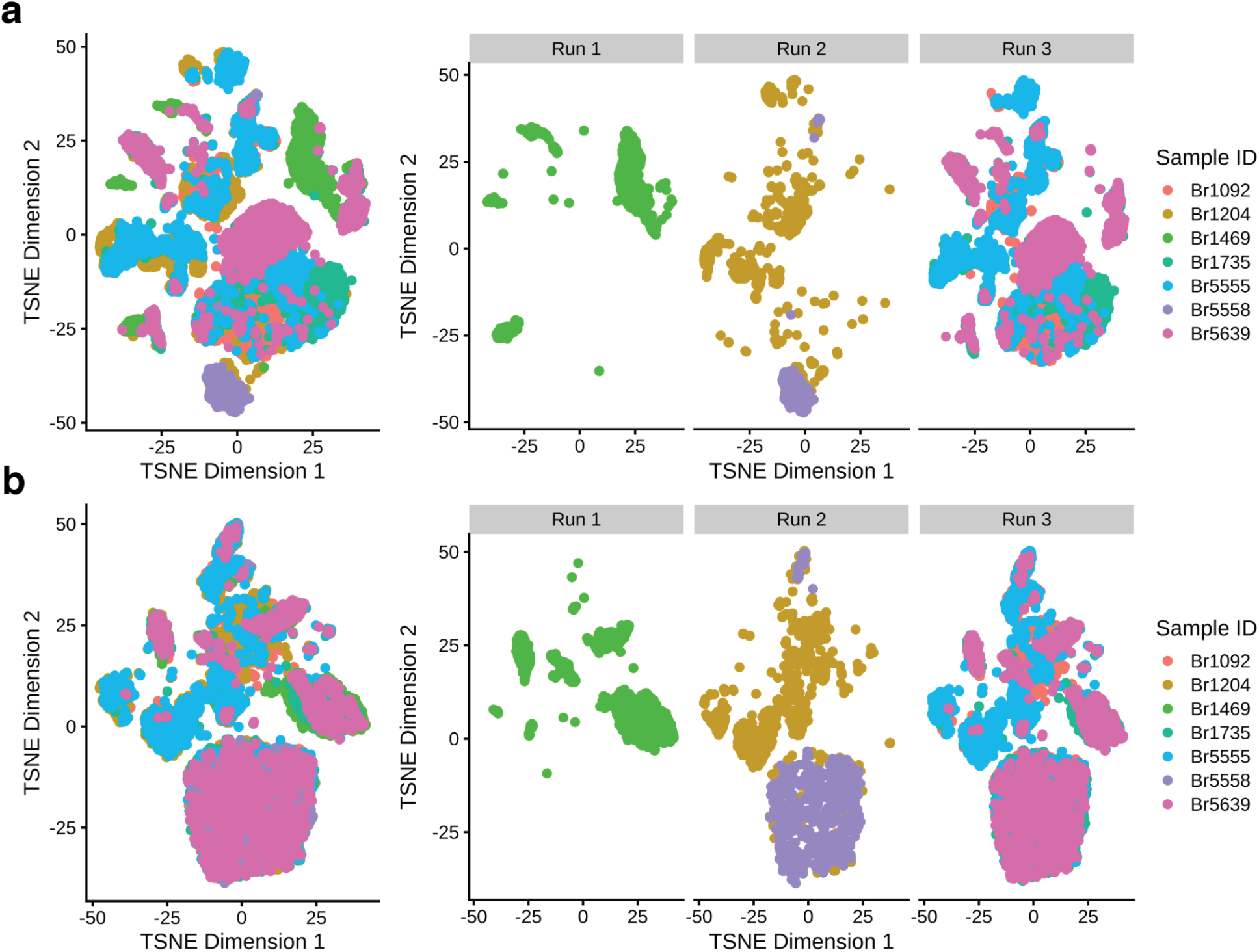
Pre and post harmony by sequencing run. Batch correction of snRNA-seq data visualized by t-SNE plots. **A**) t-SNE of principal components (PCs) pre-batch correction, colored by sample and faceted by sequencing run. **B**) t-SNE of PCs post-batch correction with *Harmony*, colored by sample and faceted by sequencing run. Batch correction with *Harmony* reduced sample and sequencing run batch effects.

**Fig. S5.**
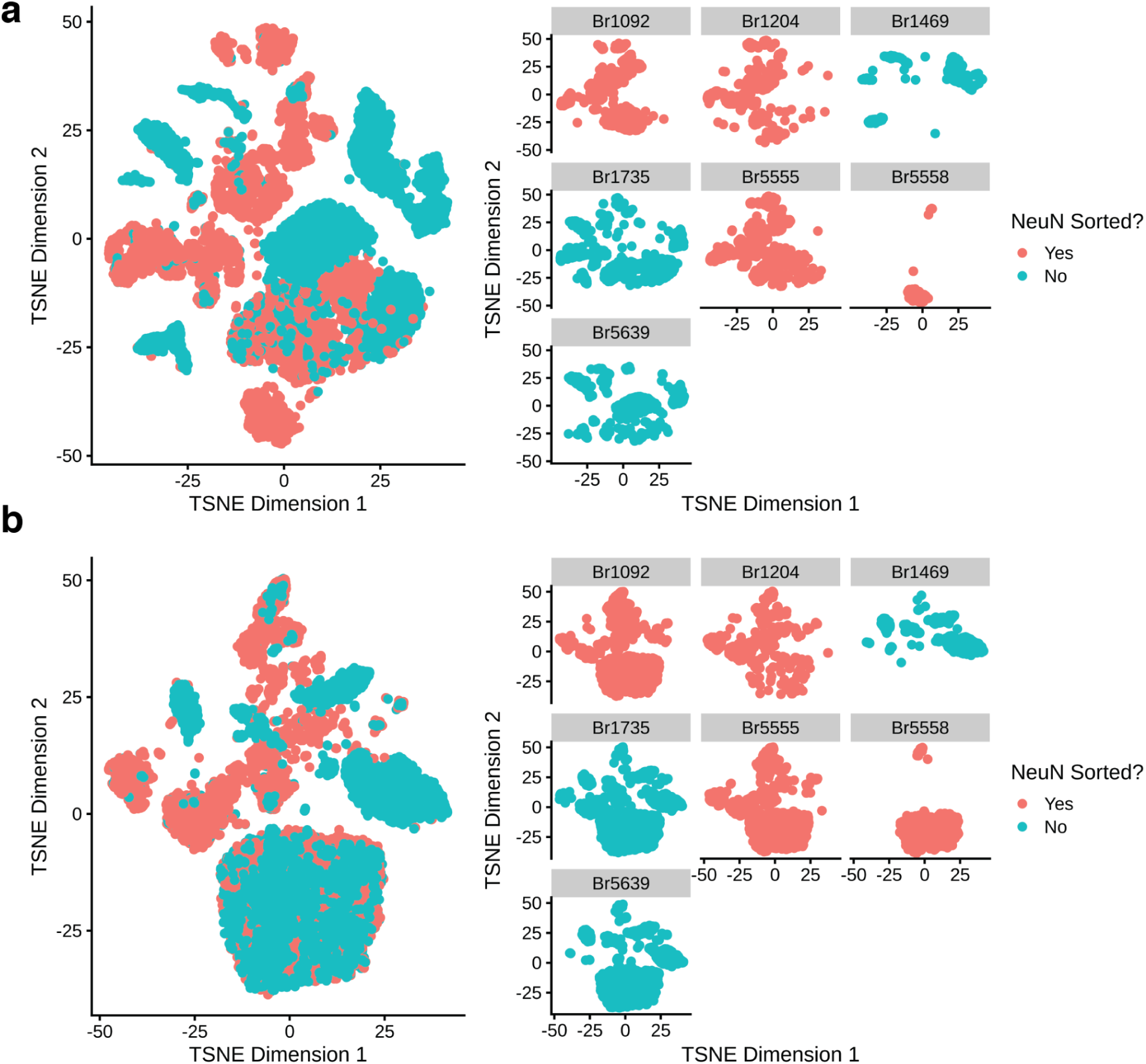
Pre and post harmony by NeuN sorting. Batch correction of snRNA-seq data visualized by t-SNE plots. **A**) t-SNE of principal components (PCs) pre-batch correction, colored by neuronal enrichment performed with NeuN antibody labeling and FANS. **B**) t-SNE of PCs post-batch correction with *Harmony*, colored by neuronal enrichment performed with NeuN antibody labeling and FANS. While batch correction with *Harmony* reduced NeuN sorting differences, some differences remain. This is expected given that the NeuN sorted samples are enriched for neuronal cell types compared to samples without NeuN sorting.

**Fig. S6.**
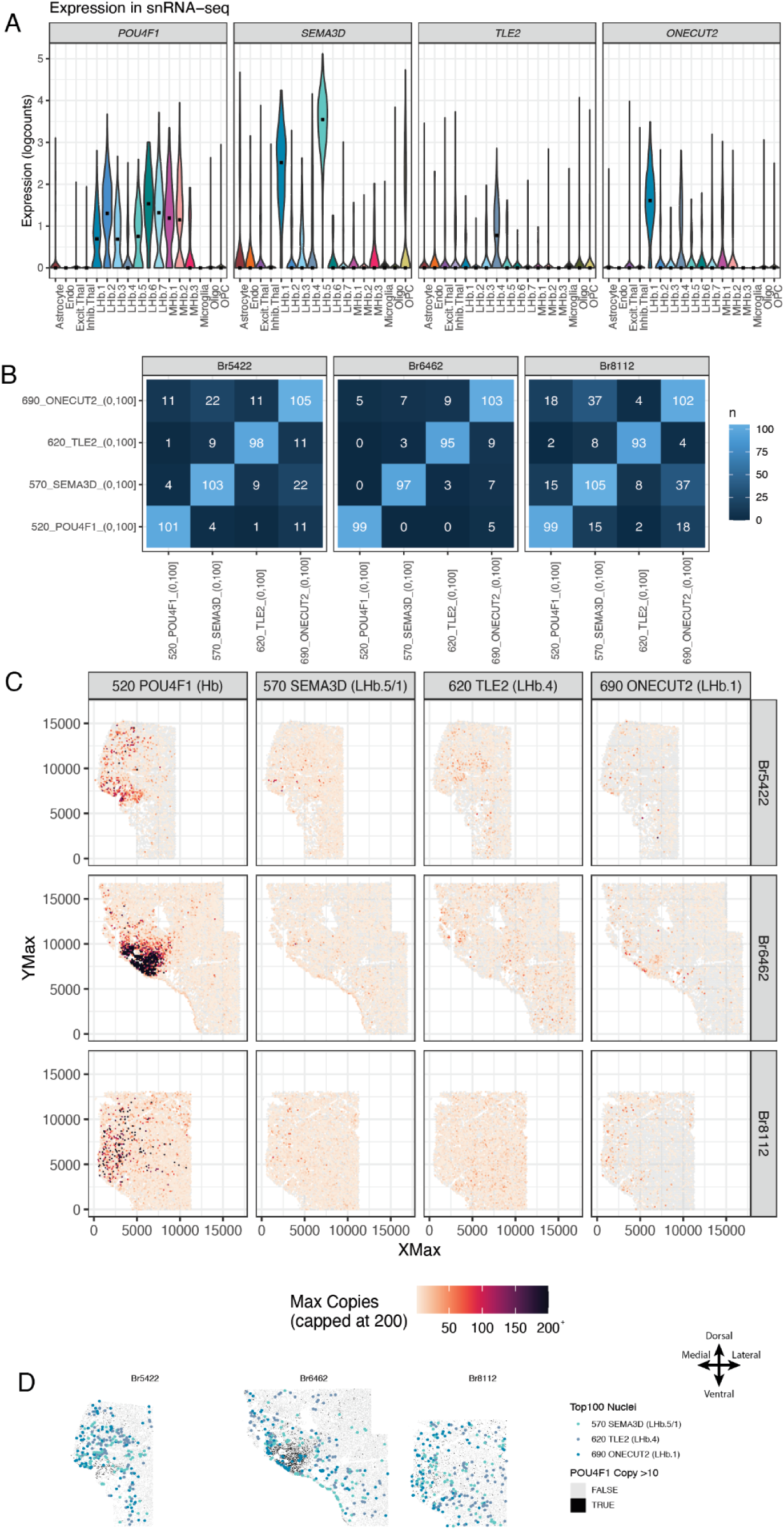
Lateral habenula smFISH to visualize LHb.1, LHb.4, and LHb.5 subpopulations. **A**) Violin plots of selected mean ratio marker gene expression (in normalized (`logcounts`)) for *POU4F1, SEMA3D, TLE2, ONECUT2* across cell types identified from the snRNA-seq data. *POU4F1* broadly marks Hb cell populations (labeled with Opal dye 520)*, SEMA3D* marks LHb.1/5 (labeled with Opal dye 570)*, TLE2* marks LHb.4 (labeled with Opal dye 620), and *ONECUT2* marks LHb.1 (labeled with Opal dye 690). **B**) For each donor tissue section: Confusion matrix of top 100 cells per marker gene probe channel, ranked by number of transcript copies. Non-diagonal squares (dark blue) show number of cells that are in the top 100 of those two marker genes. Ranks include ties, therefore each square along the diagonal (light blue) can be more than 100. **C**) For each marker gene probe channel: Hexbin plots displaying the maximum number of transcript copies per bin across the tissue section from each donor. To aid visualization, all values > 200 copies are the same color. XMax and YMax values provide spatial locations of the cells defined during cell object segmentation in HALO. **D**) Spatial plots displaying the top cells expressing each marker gene (**Fig. 3Ai**), with each detected cell in the tissue section plotted as a gray or black rectangle. Cells in black with > 10 transcript copies of the Hb marker gene *POU4F1* depict the habenula region. Colored circles mark the spatial location of the top 100 ranked cells that most robustly express each marker gene probe. If a cell was ranked in the top 100 for more than one marker gene (see **Fig. S6B**), it was colored by the gene for which it had the most number of transcript copies.

**Fig. S7.**
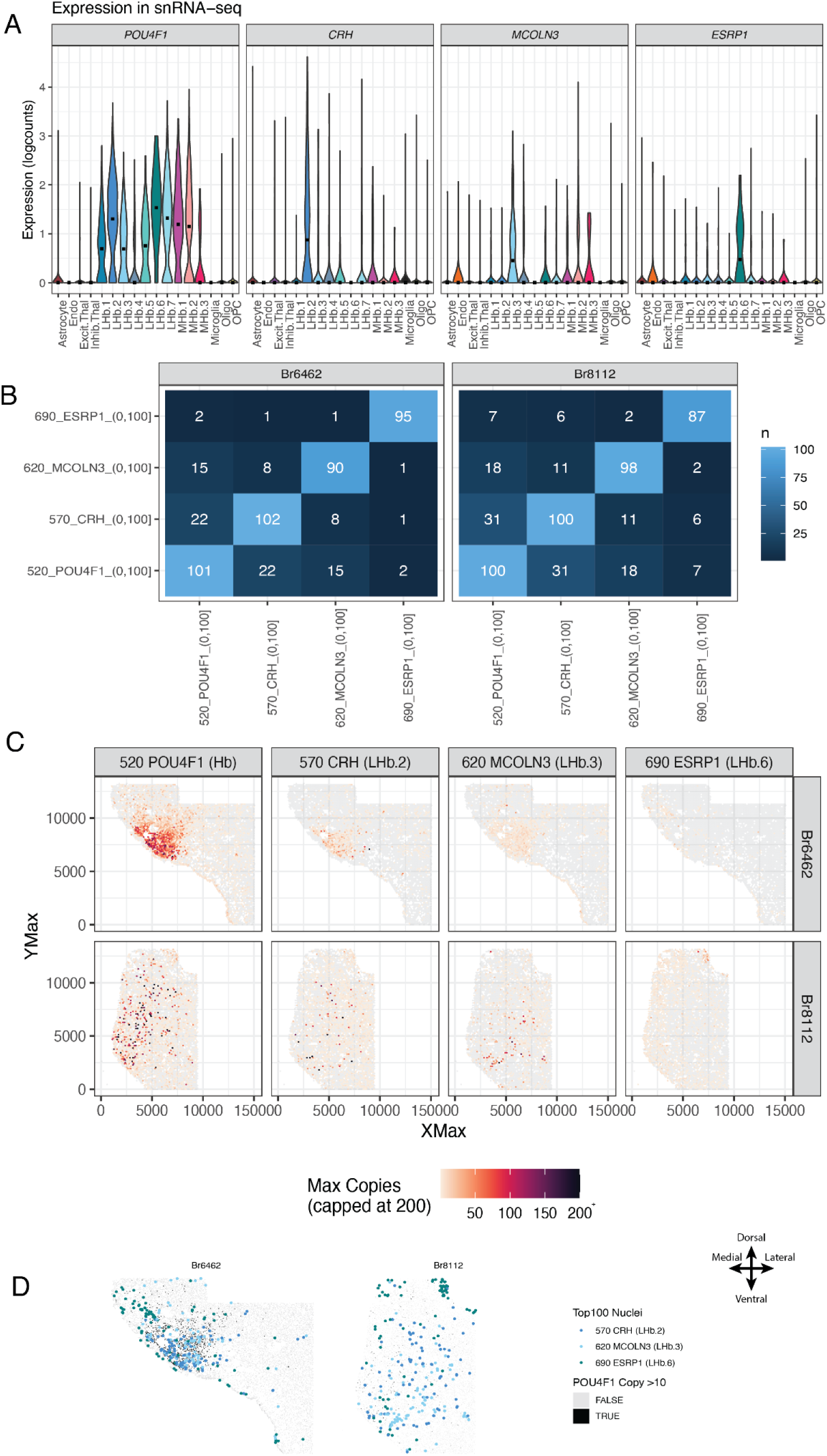
Lateral habenula smFISH to visualize LHb.2, LHb.3, and LHb.6 subpopulations. **A)** Violin plots of selected mean ratio marker gene expression (in normalized (`logcounts`)) for *POU4F1, CRH, MCOLN3, ESRP1* across cell types identified from the snRNA-seq data. *POU4F1* broadly marks Hb cell populations (labeled with Opal dye 520)*, CRH* marks LHb.2 (labeled with Opal dye 570)*, MCOLN3* marks LHb.3 (labeled with Opal dye 620), and *ESRP1* marks LHb.6 (labeled with Opal dye 690). **B)** For each donor tissue section: Confusion matrix of top 100 cells per marker gene probe channel, ranked by number of transcript copies. Non-diagonal squares (dark blue) show number of cells that are in the top 100 of those two marker genes. Ranks include ties, therefore each square along the diagonal (light blue) can be more than 100. **C)** For each marker gene probe channel: Hexbin plots displaying the maximum number of transcript copies per bin across the tissue section from each donor. To aid visualization, all values > 200 copies are the same color. XMax and YMax values provide spatial locations of the cells defined during cell object segmentation in HALO. **D**) Spatial plots displaying the top cells expressing each marker gene (**Fig. 3Bi**), with each detected cell in the tissue section plotted as a gray or black rectangle. Cells in black with > 10 transcript copies of the Hb marker gene *POU4F1* depict the habenula region. Colored circles mark the spatial location of the top 100 ranked cells that most robustly express each marker gene probe. If a cell was ranked in the top 100 for more than one marker gene (see **Fig. S7B**), it was colored by the gene for which it had the most number of transcript copies.

**Fig. S8.**
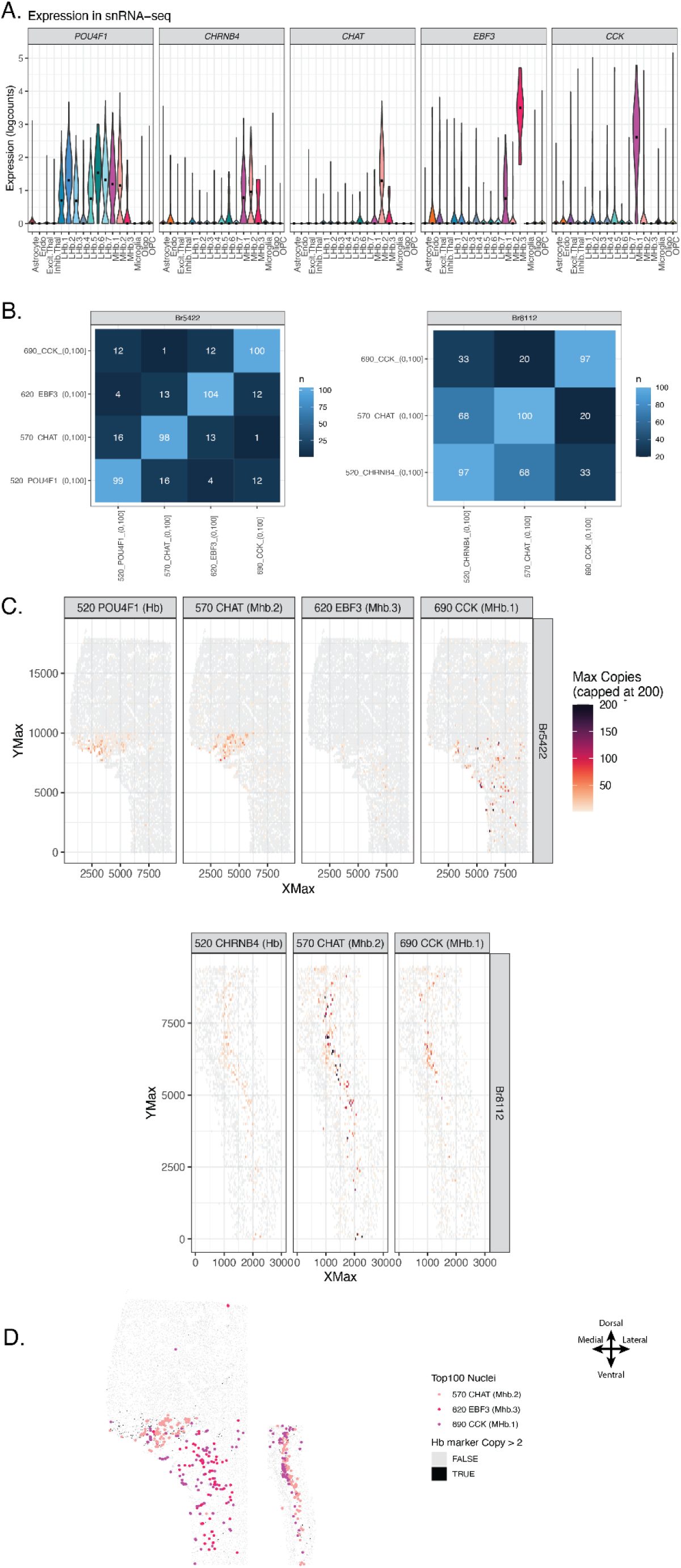
Medial habenula smFISH for MHb.1, MHb.2, and MHb.3 subpopulations. **A)** Violin plots of selected mean ratio marker gene expression (in normalized (`logcounts`)) for *POU4F1, CHAT, EBF3, CCK* across cell types identified from the snRNA-seq data. *POU4F1* broadly marks Hb cell populations (labeled with Opal dye 520)*, CHAT* marks MHb.2 (labeled with Opal dye 570)*, EBF3* marks MHb.3 (labeled with Opal dye 620), and *CCK* marks MHb.1 (labeled with Opal dye 690). **B)** For each donor tissue section: Confusion matrix of top 100 cells per marker gene probe channel, ranked by number of transcript copies. Non-diagonal squares (dark blue) show number of cells that are in the top 100 of those two marker genes. Ranks include ties, therefore each square along the diagonal (light blue) can be more than 100. **C)** For each marker gene probe channel: Hexbin plots displaying the maximum number of transcript copies per bin across the tissue section from each donor. To aid visualization, all values > 200 copies are the same color. XMax and YMax values provide spatial locations of the cells defined during cell object segmentation in HALO. **D**) Spatial plots displaying the top cells expressing each marker gene (**Fig. 3Ci**), with each detected cell in the tissue section plotted as a gray or black rectangle. Cells in black with > 2 transcript copies of the Hb marker gene *POU4F1* depict the habenula region. Colored circles mark the spatial location of the top 100 ranked cells that most robustly express each marker gene probe. If a cell was ranked in the top 100 for more than one marker gene (see **Fig. S8B**), it was colored by the gene for which it had the most number of transcript copies.

**Fig. S9.**
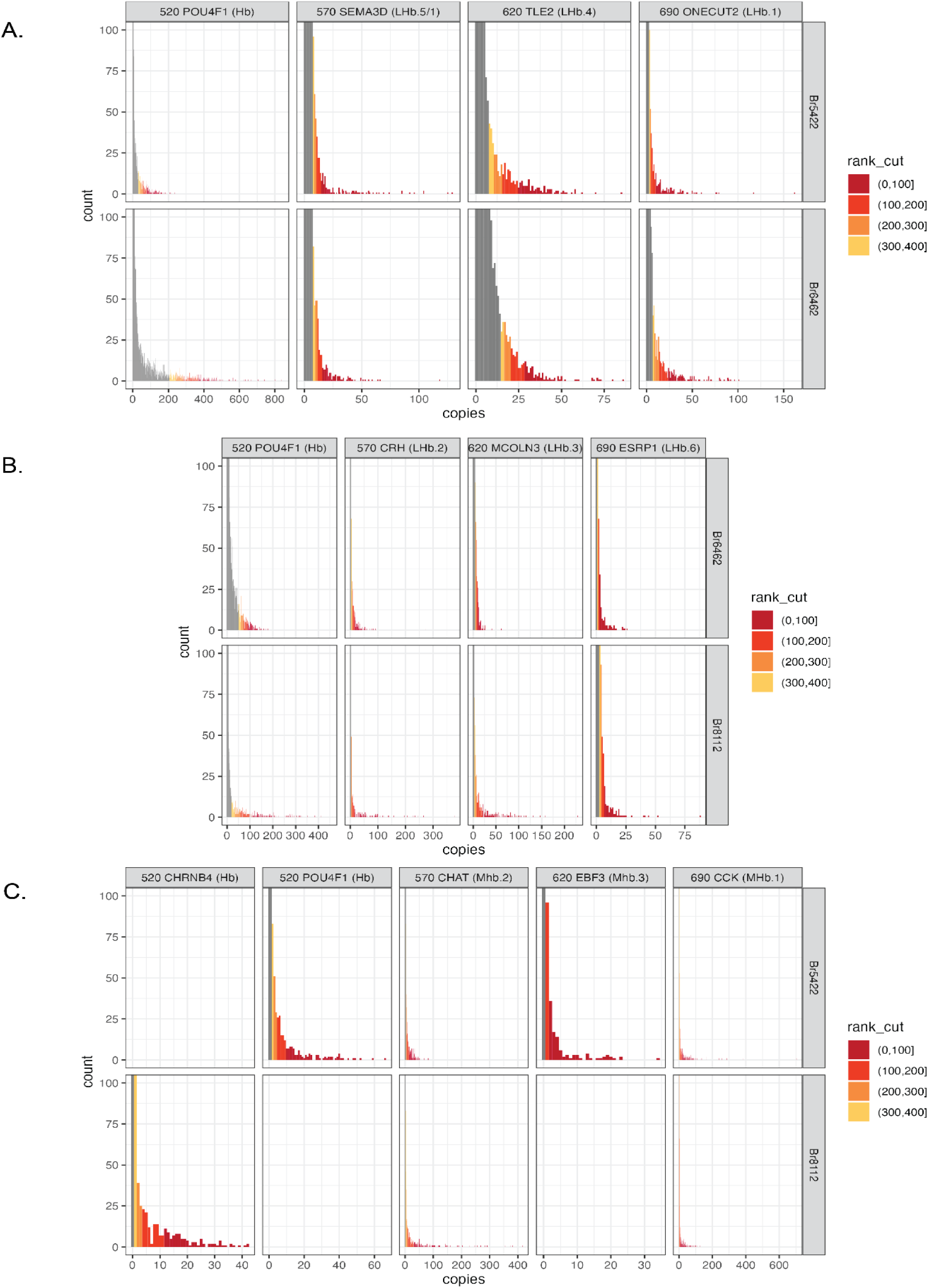
Distributions of marker gene expression estimations across smFISH experiments. HALO software (Indica Labs) was used to detect and quantify RNAScope probes in each cell object. **A)** Frequency count distribution of the estimated number of *POU4F1*, *SEMA3D*, *TLE2*, and *ONECUT2* marker gene copies for each cell object. **B)** Frequency count distribution of the estimated number of *POU4F1*, *CRH*, *MCOLN3*, and *ESRP1* marker gene copies for each cell object. **C)** Frequency count distribution of the estimated number of *POU4F1*, *CHAT*, *EBF3*, *CCK*, and *CHRNB4* marker gene copies for each cell object. In A-C, the count frequencies (y-axis) are limited to 100 in order to observe the frequency counts for non-zero copy values.

**Fig. S10.**
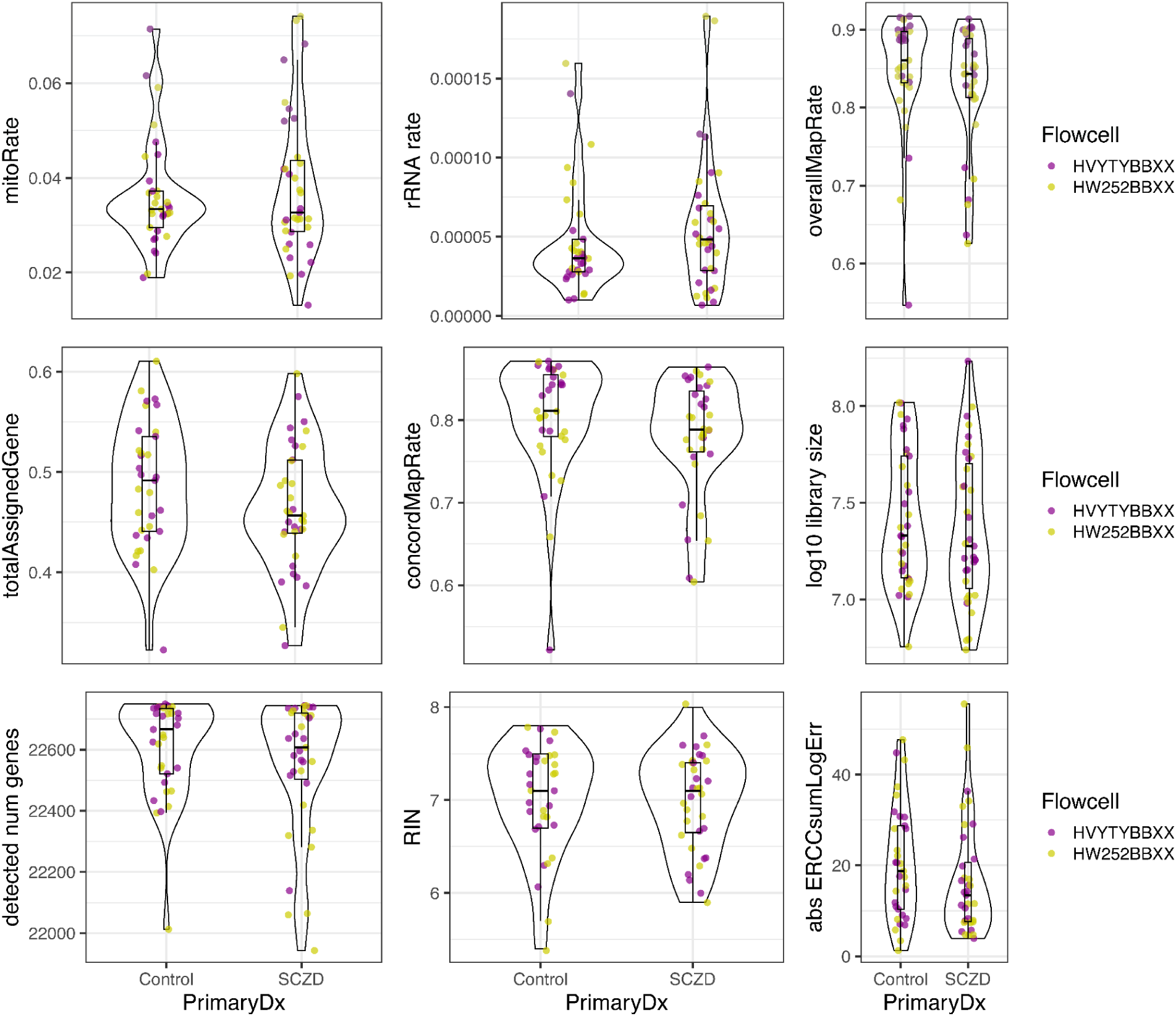
Bulk RNA-sequencing quality control metrics. Boxplots and violin distribution graphs for several bulk RNA-seq quality metrics computed by *SPEAQeasy*. Samples are separated by Control vs. SCZD status and colored by the sequencing flowcell. Sample Br5572, which was excluded from analysis, is not included in these graphs.

**Fig. S11.**
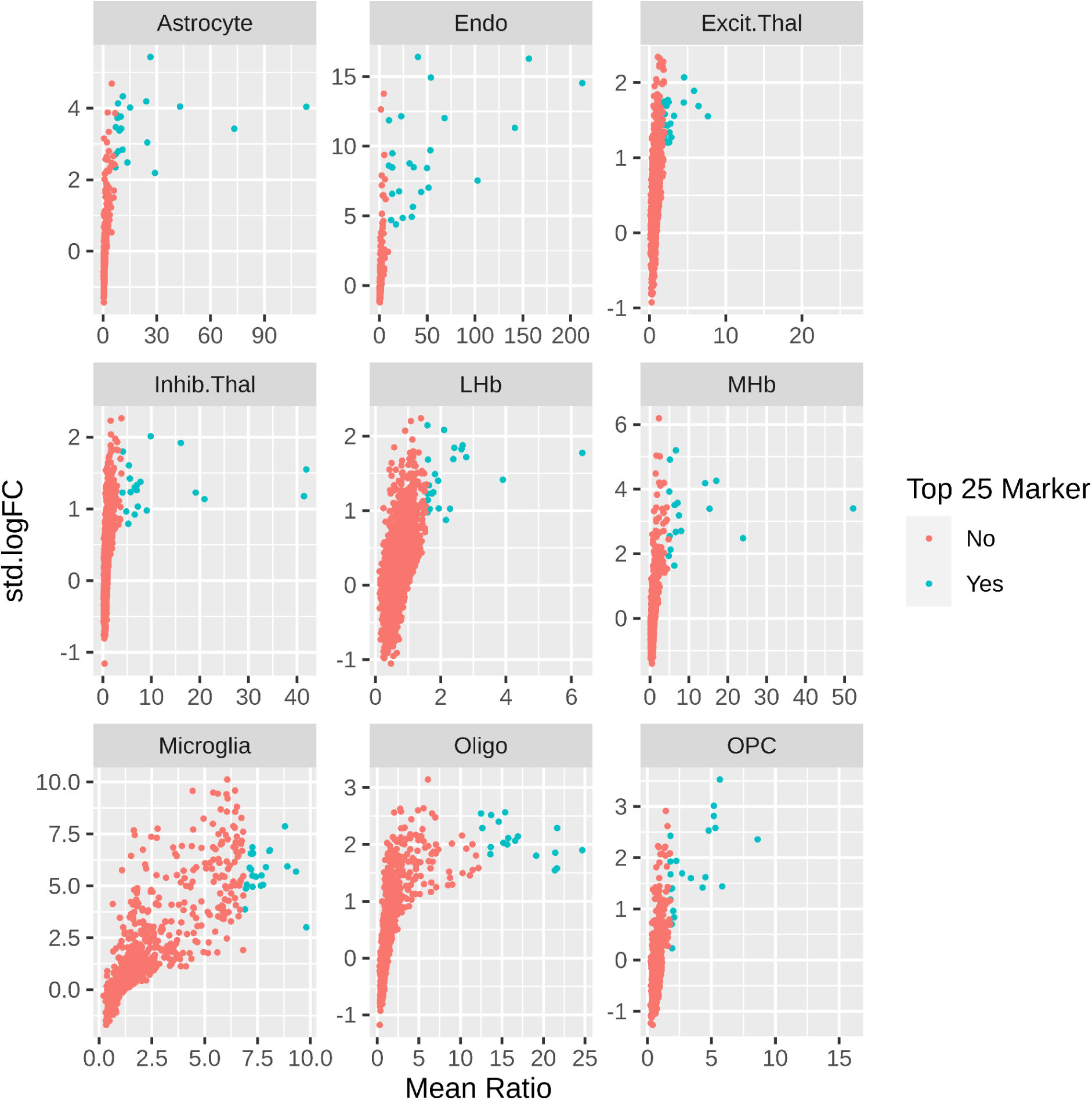
Selection of top mean ratio marker genes for cell type deconvolution. For each cell type category, mean ratios for all the detected genes (see **Supplemental Methods**) were computed by `getMeanRatio2()` from *DeconvoBuddies* and compared against the genes’ standard log2 fold change values (std.logFC), which were computed by `findMarkers()` from *scran*. The top 25 mean ratio marker genes are shown in teal, and typically have a high standard log2 fold change value.

**Fig. S12.**
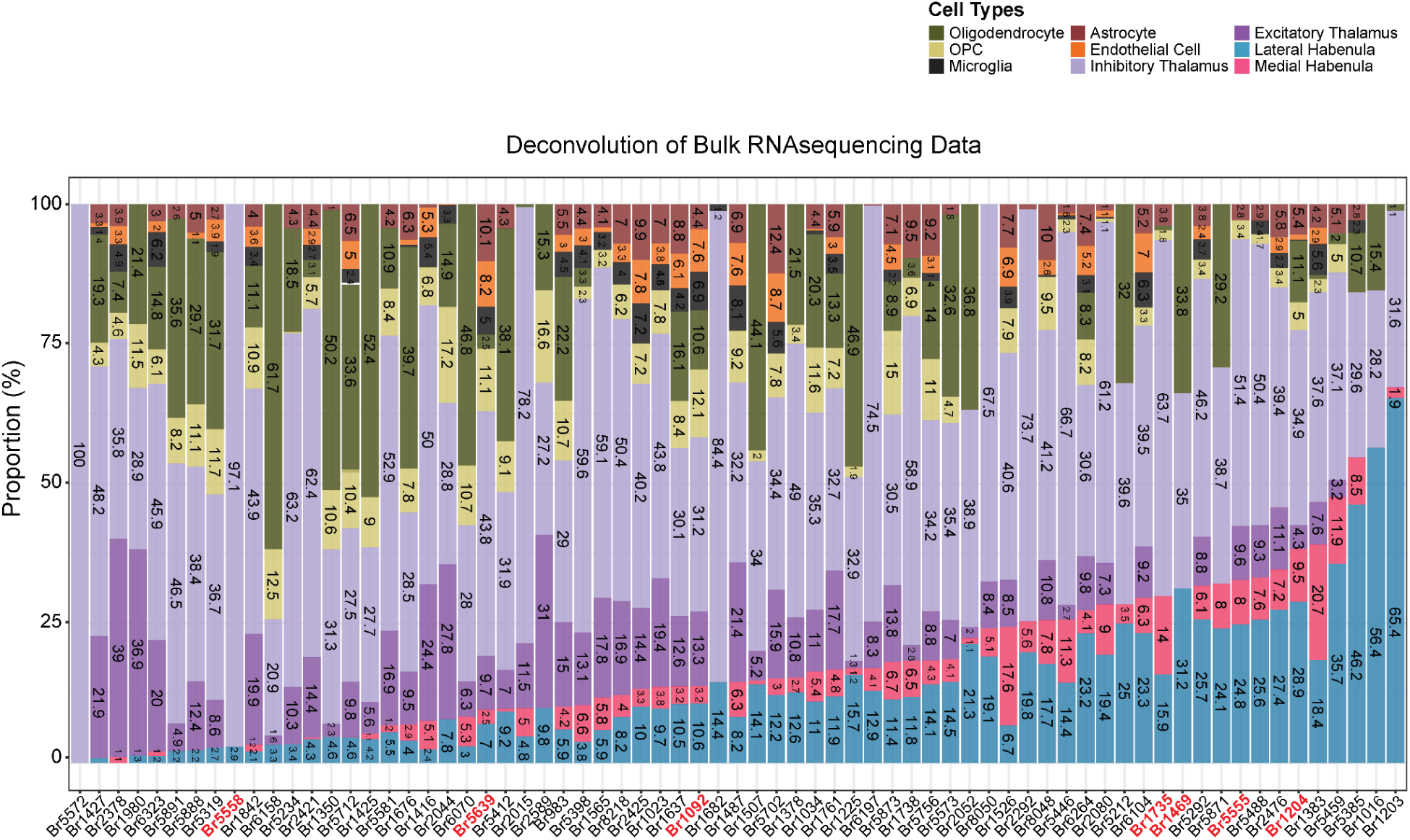
Cell type proportions of the bulk RNA-sequencing samples as estimated by deconvolution. Bar plots show the proportions of each cell type in the bulk RNA-seq samples as estimated by deconvolution. Increasing from left to right, samples are ordered by the sum of their estimated LHb (blue) and MHb (pink) proportions. As sample Br5572 was estimated to consist entirely of inhibitory thalamic neurons, we dropped this sample from further analyses. Following this exclusion, we retained 33 Control and 35 SCZD samples. Subjects highlighted in red were also used for snRNA-seq.

**Fig. S13.**
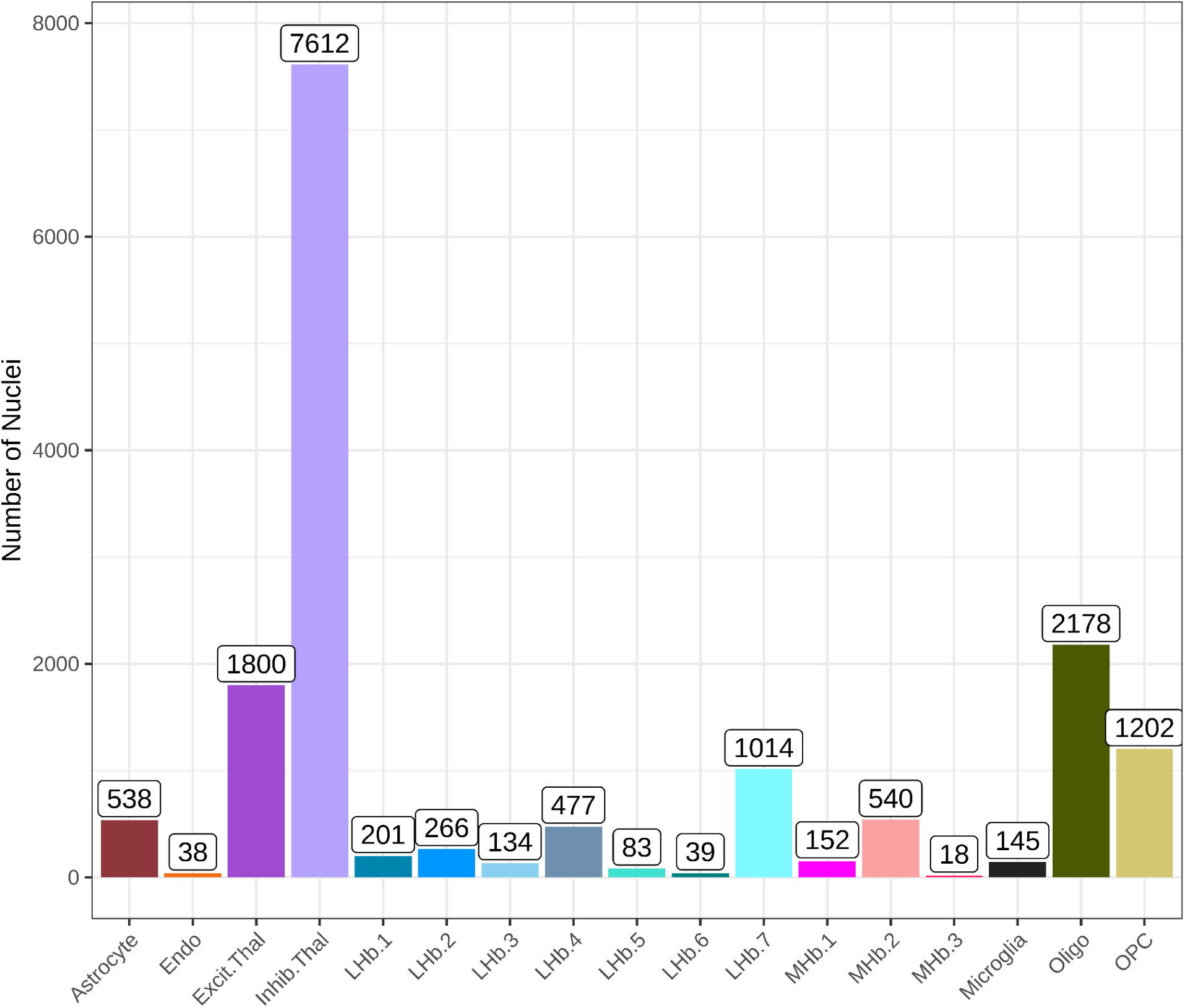
snRNA-seq post QC nuclei counts by cell type. Barplots showing the number of nuclei by cell type after quality control steps, aggregated across all seven snRNA-seq samples. Total nuclei post-QC: 16,437.

**Fig. S14.**
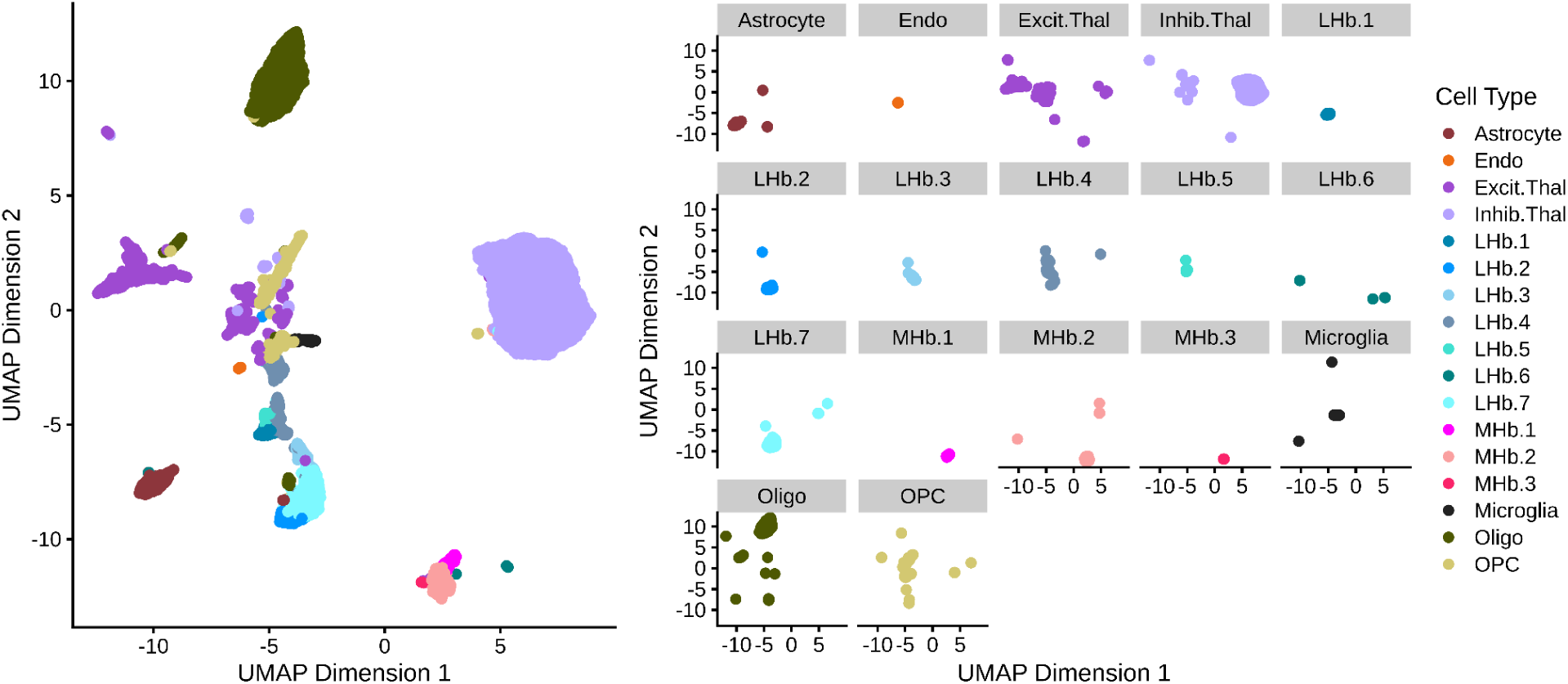
UMAP representation of the snRNA-seq data. Post-QC snRNA-seq data visualized using UMAP (similar to **Fig. 1**), colored by cell type. The right side shows the UMAP faceted by cell type.

**Fig. S15.**
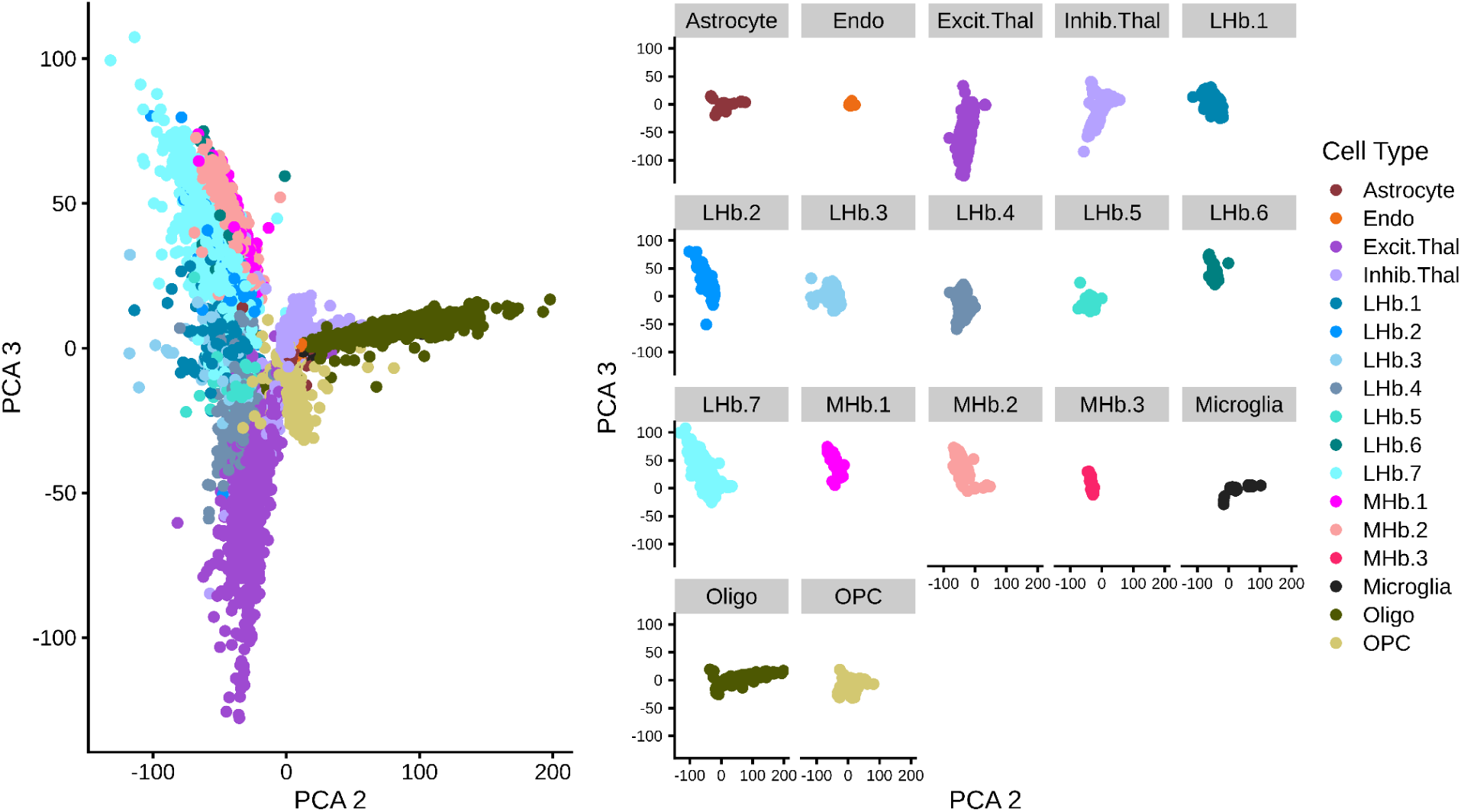
snRNA-seq principal components (PCs) by cell type. PC2 versus 3 colored by cell type clusters. The left side of the plot displays all nuclei together, while the right side shows them faceted by cell type. PC2 differentiates non-neuronal cell types (i.e. Oligodendrocytes, Microglia, Astrocytes, Endothelial cells, Oligodendrocyte Progenitor Cells) from neuronal cell types. PC3 differentiates excitatory thalamic neurons from habenula neuron subpopulations.

**Fig. S16.**
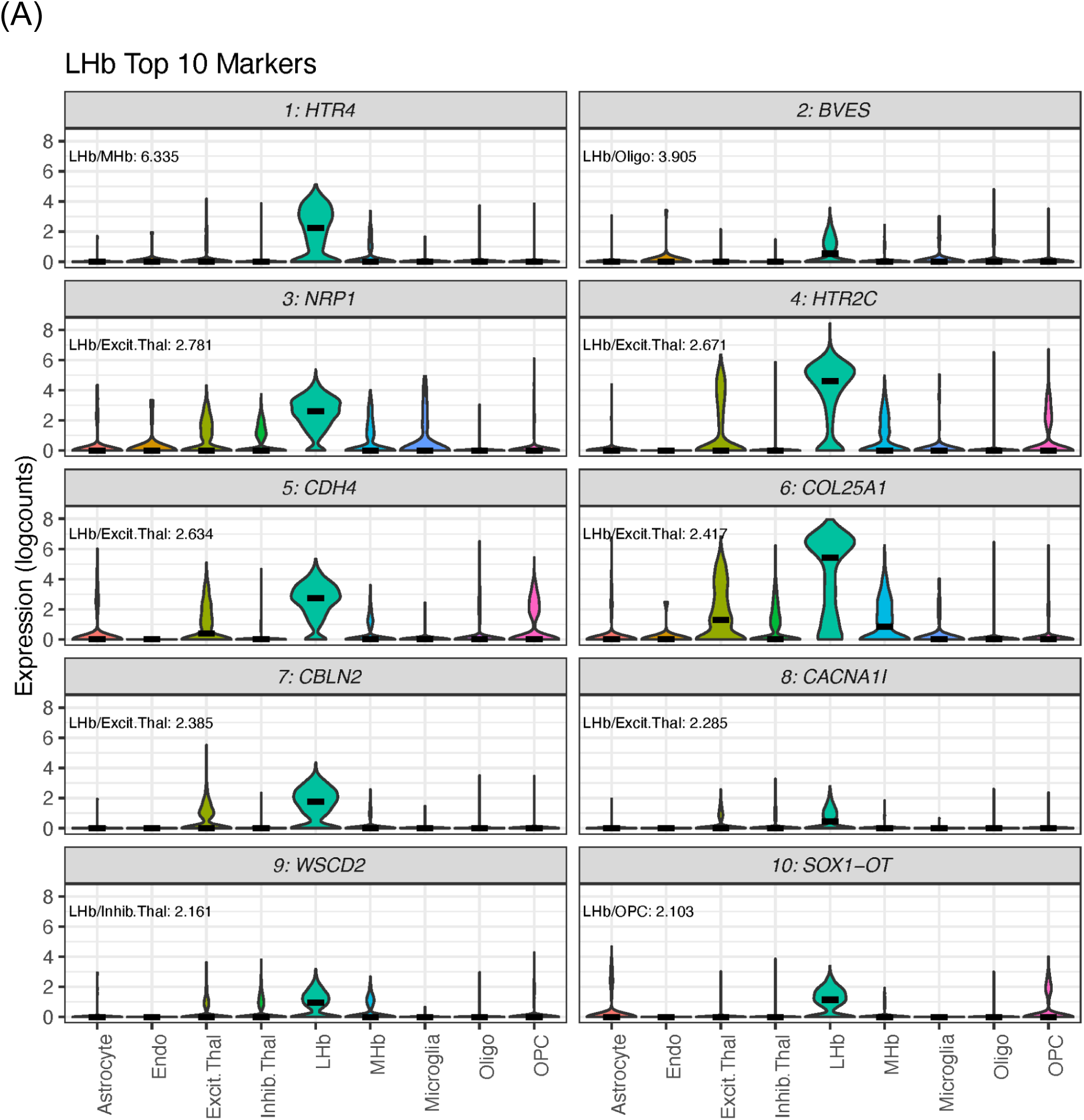
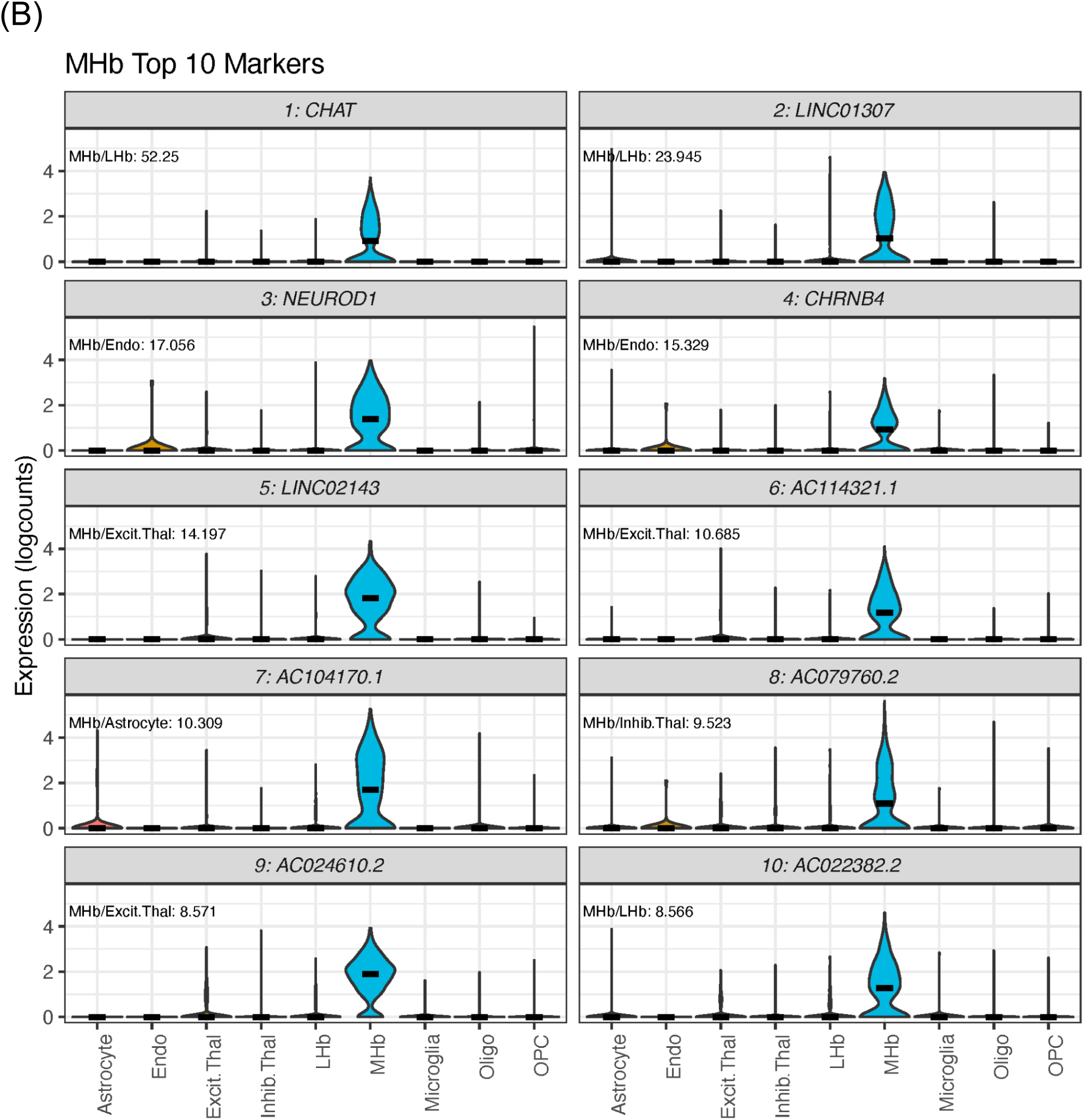
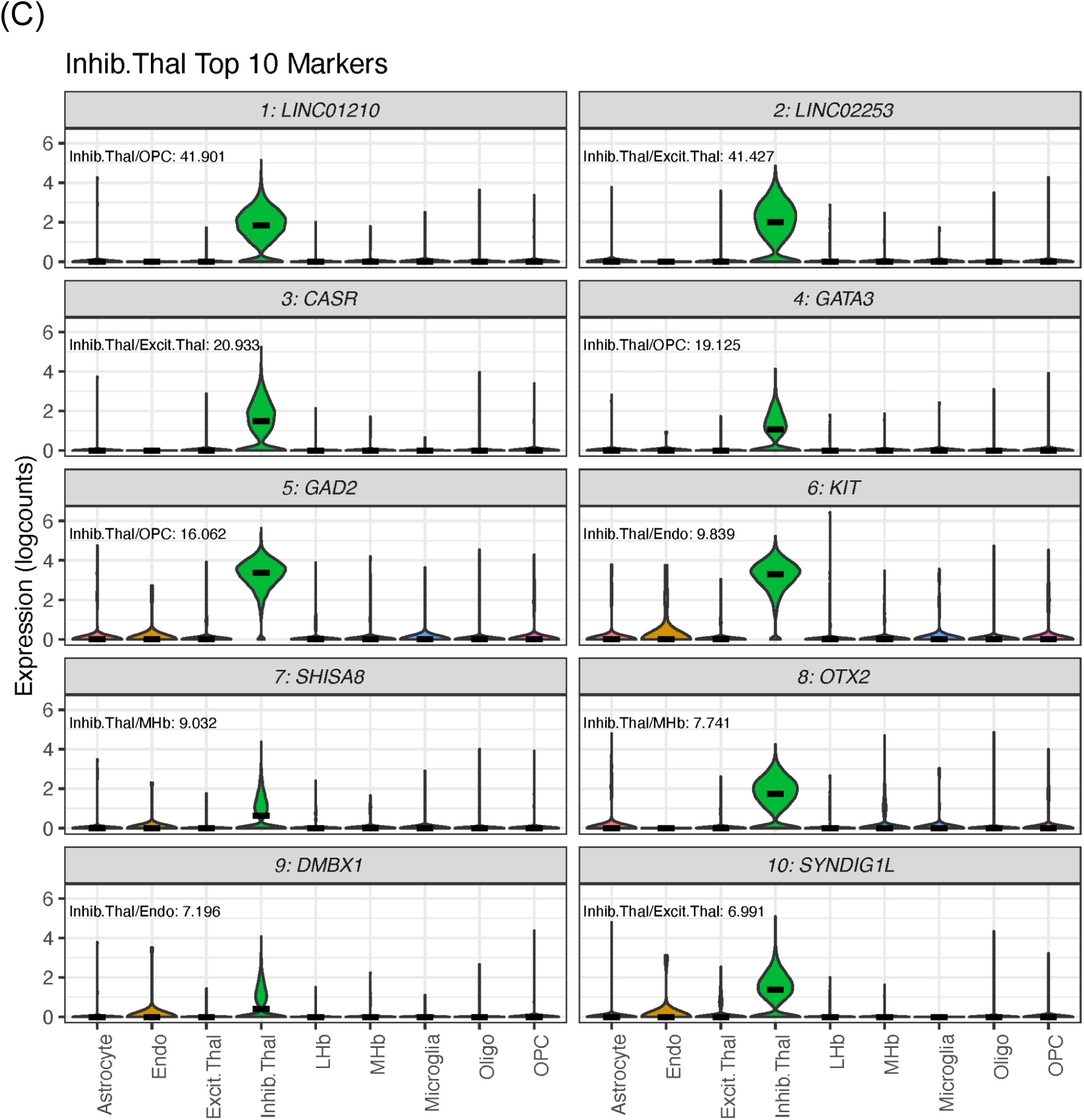
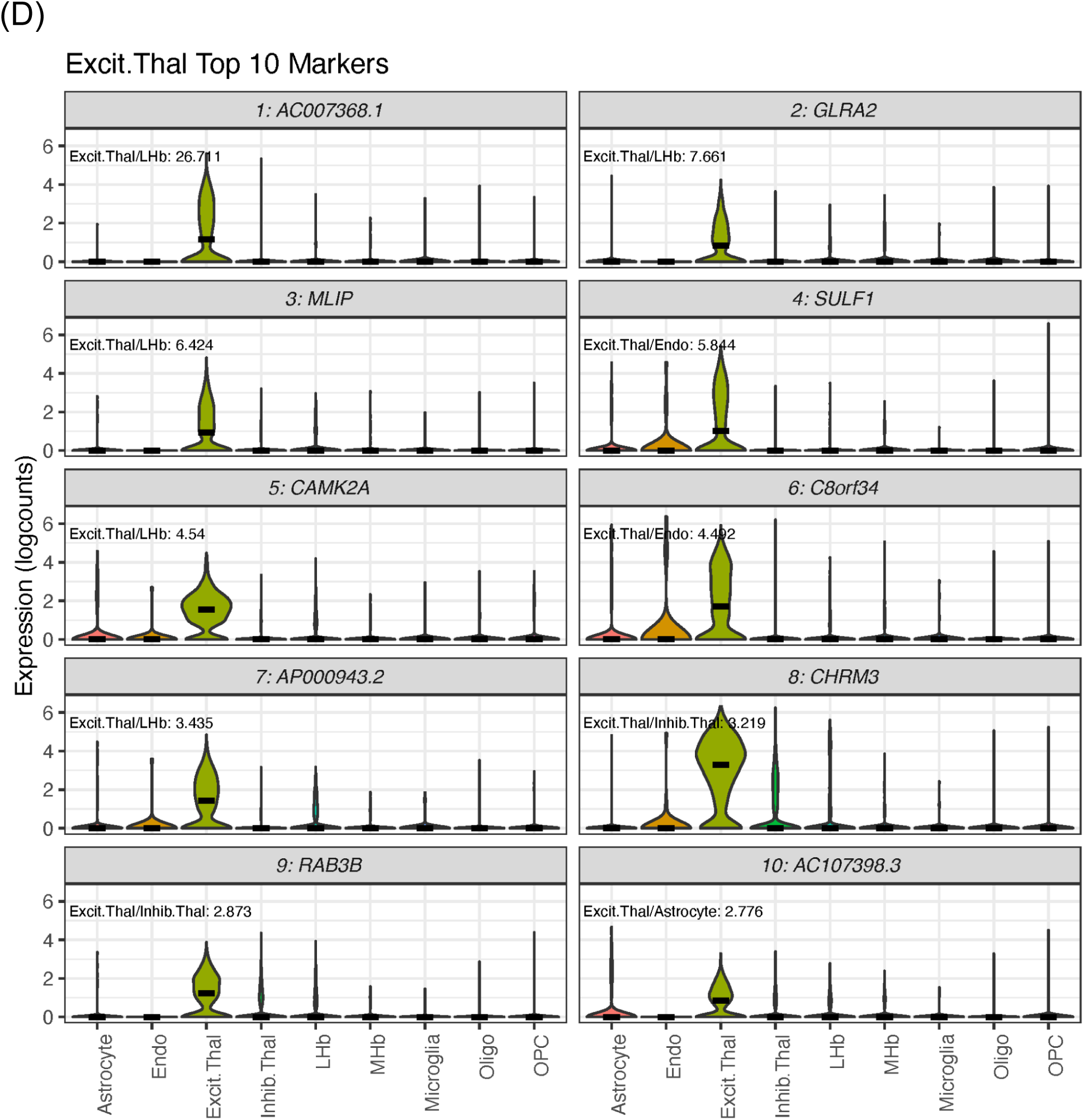
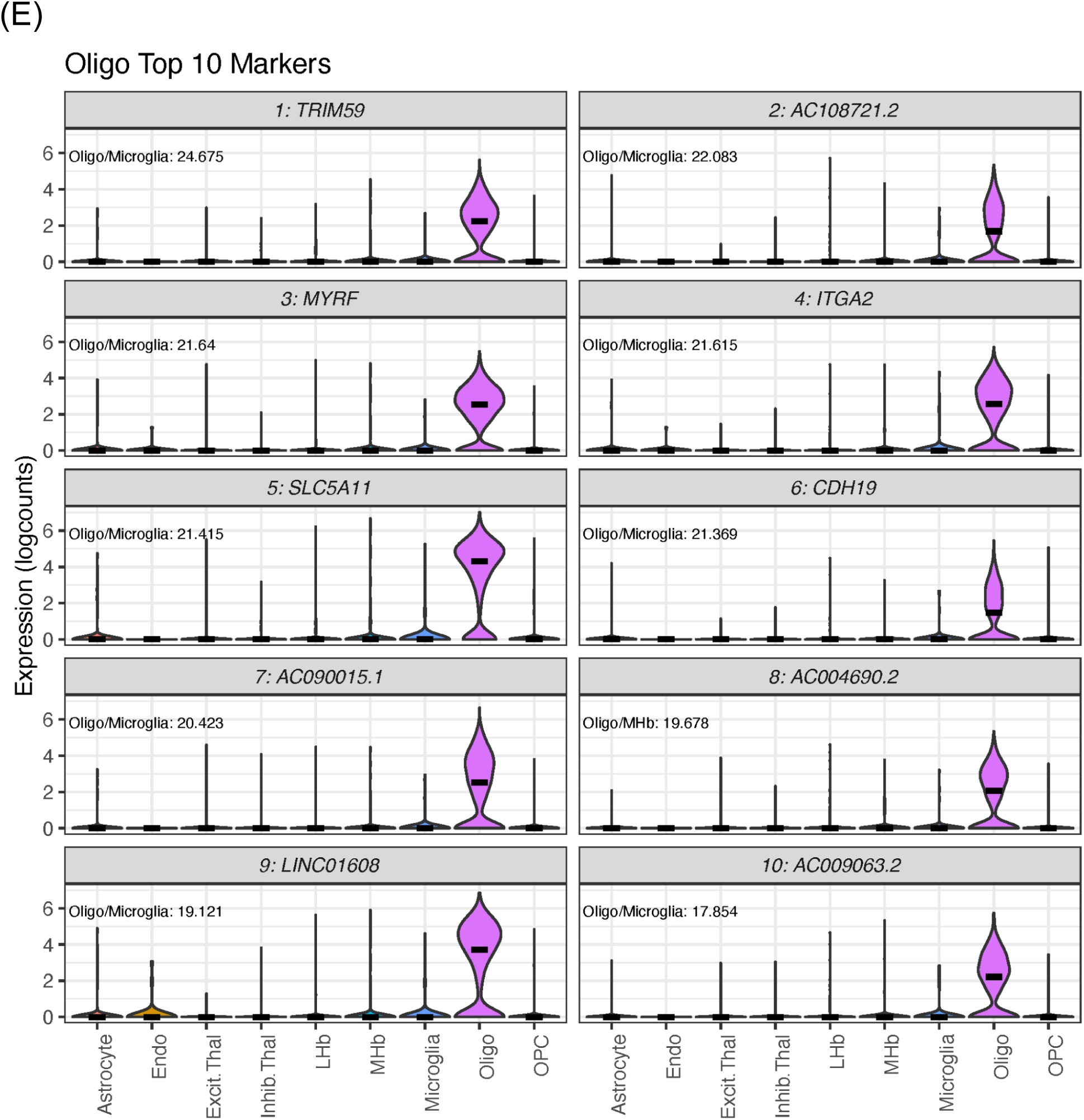
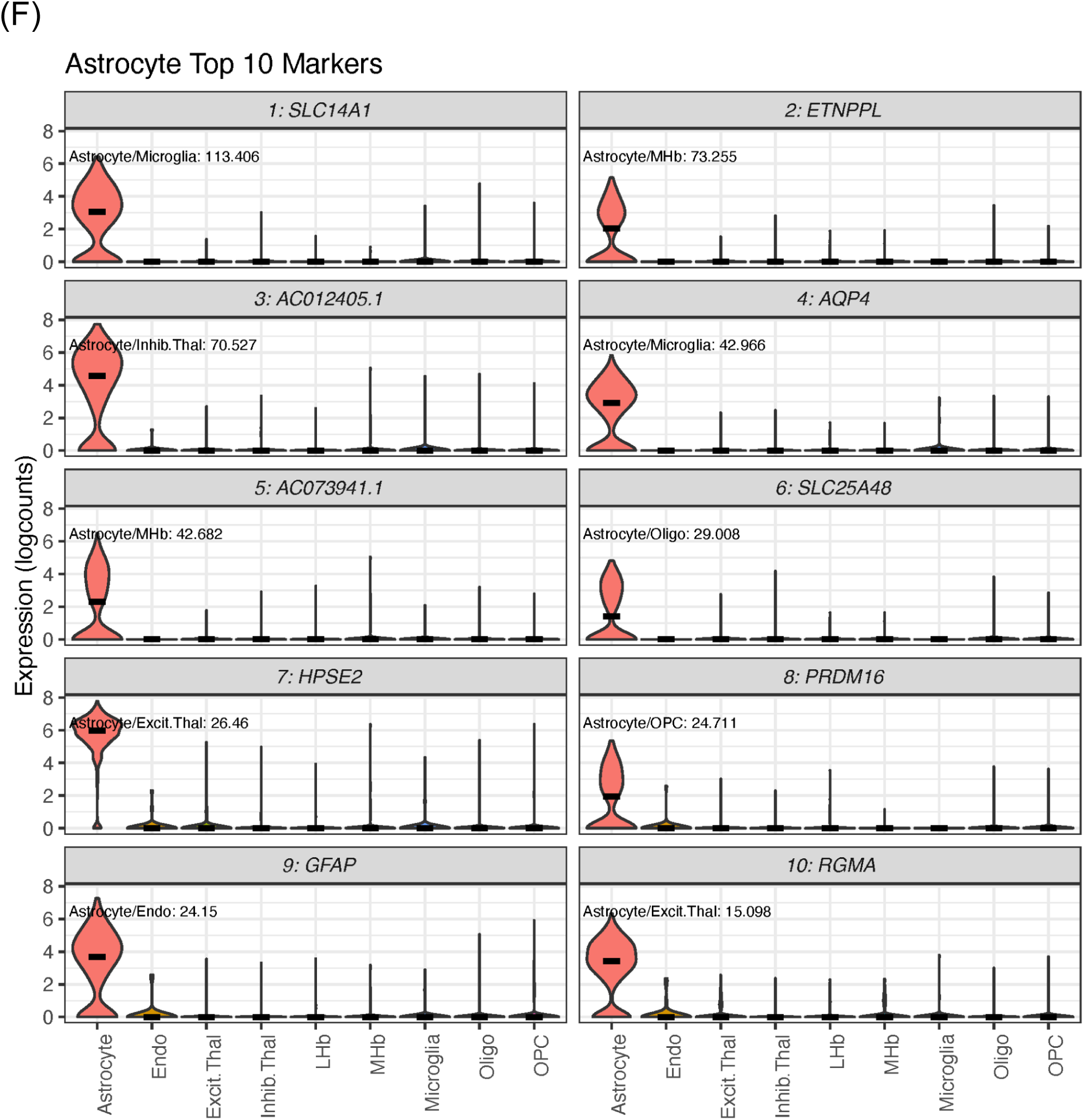
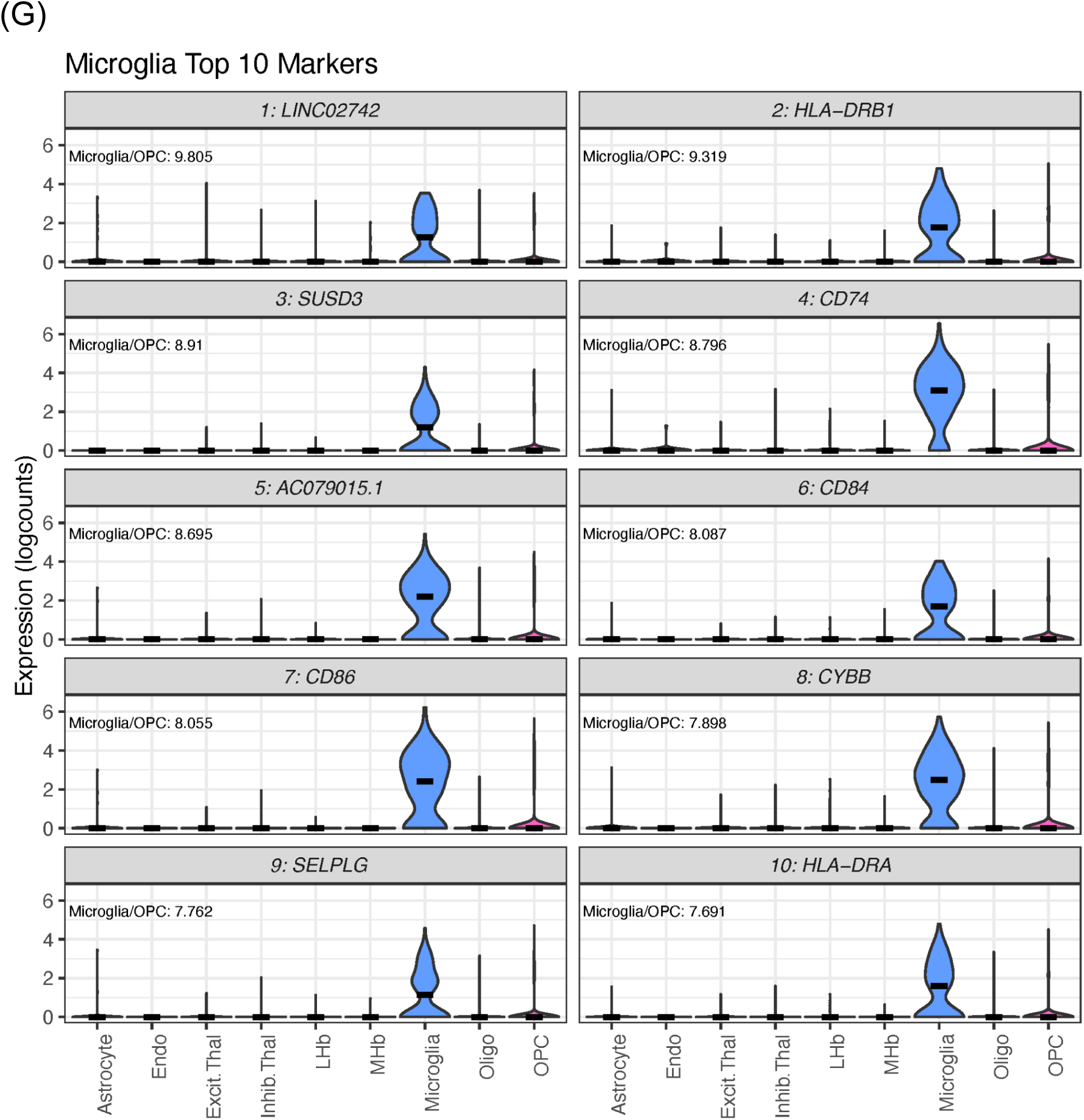
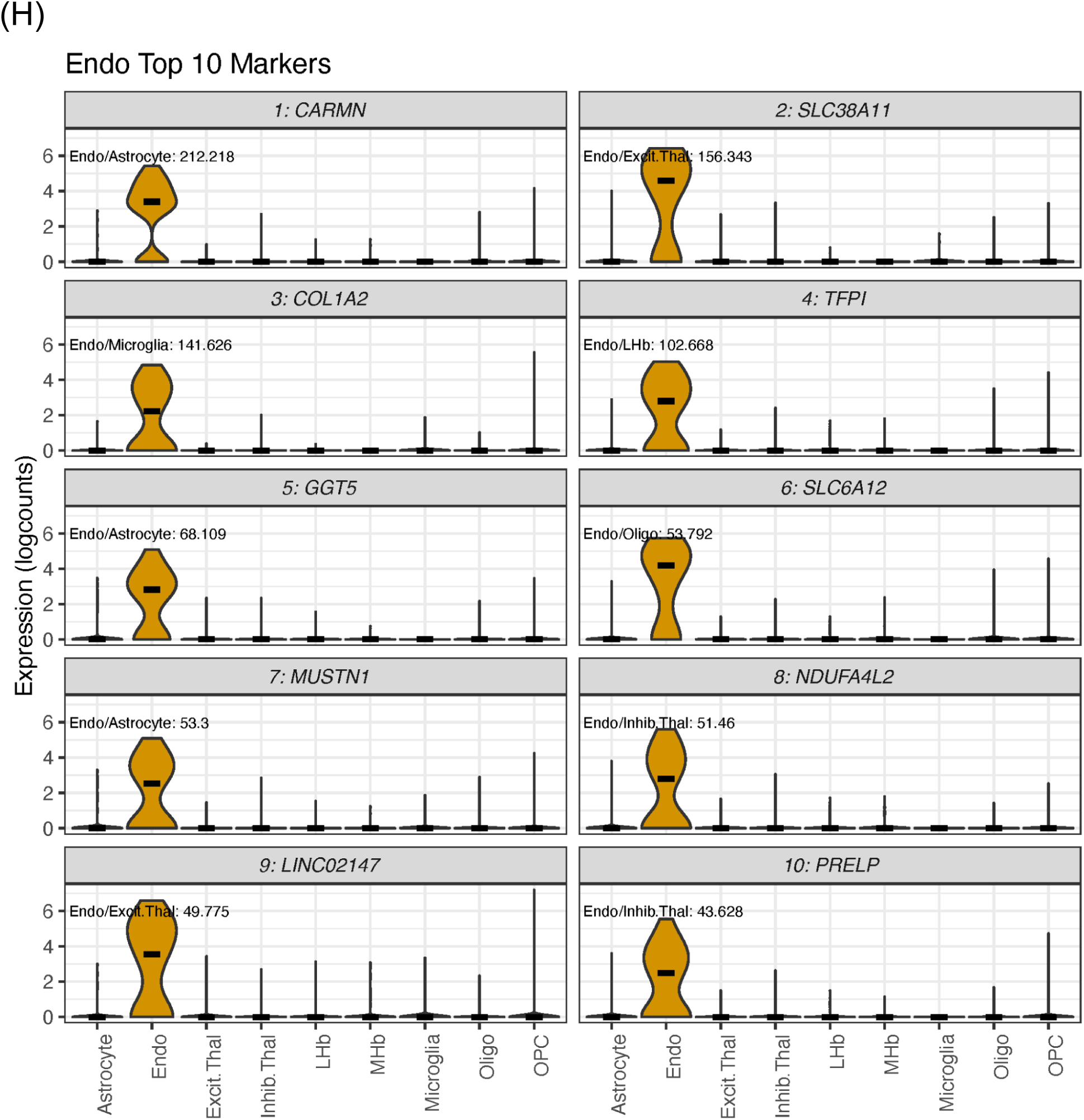
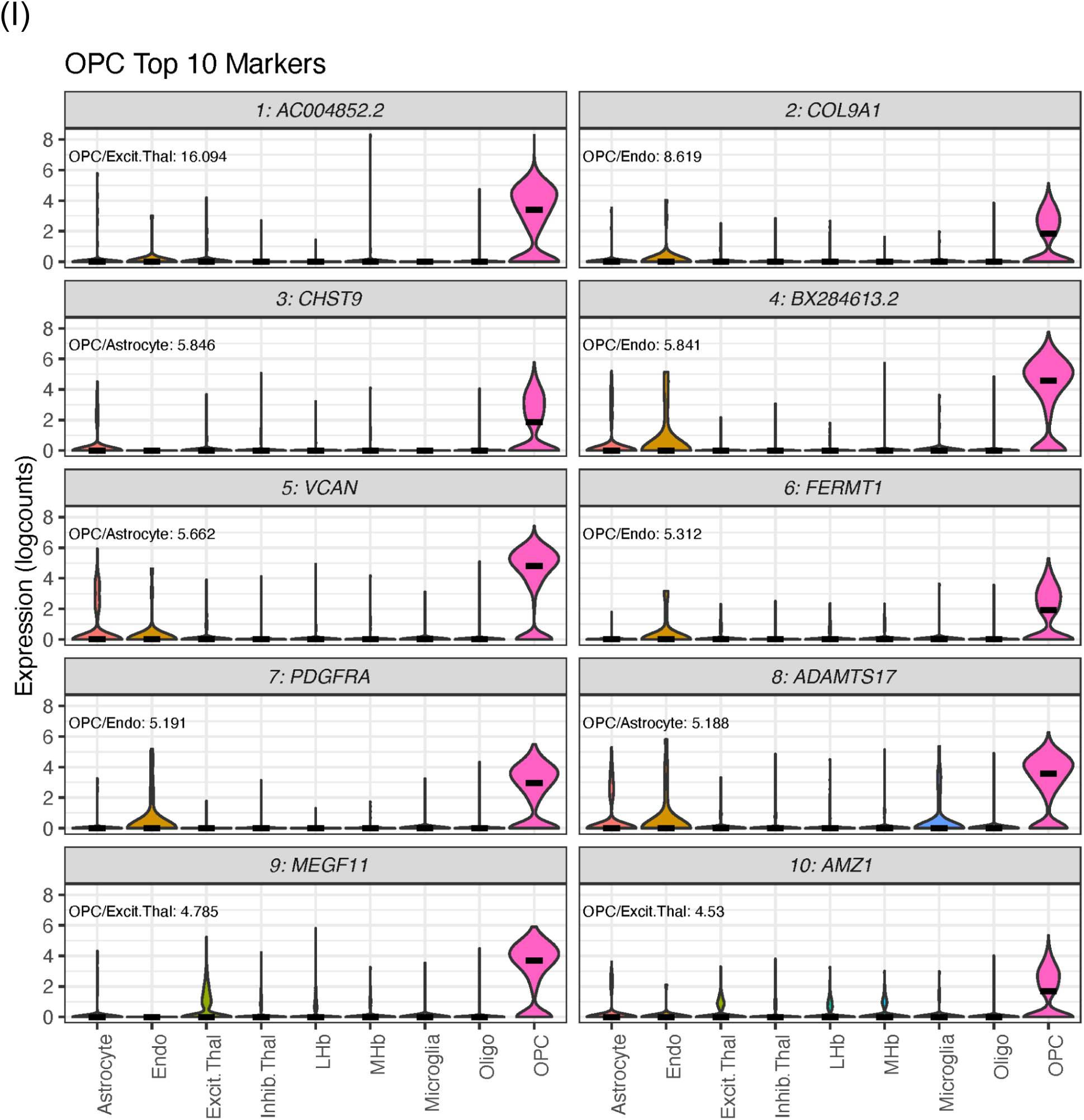
Gene expression plots for the top 10 marker genes for each cell type. Violin distribution plots of the log-normalized expression counts (logcounts) of the top 10 mean ratio marker genes for each cell type. Top left corner of each plot shows the target cell type and second-most highly expressing cell type, as well as the ratio of their mean marker gene expression levels (i.e. the mean ratio). **A**) Lateral Habenula neurons (LHb), **B**) Medial Habenula neurons (MHb), **C**) Inhibitory Thalamic neurons (Inhib.Thal), **D**) Excitatory Thalamic neurons (Excit.Thal), **E**) Oligodendrocytes (Oligo), **F**) Astrocytes, **G**) Microglia, **H**) Endothelial cells (Endo), **I**) Oligodendrocyte Progenitor Cells (OPC).

**Fig. S17.**
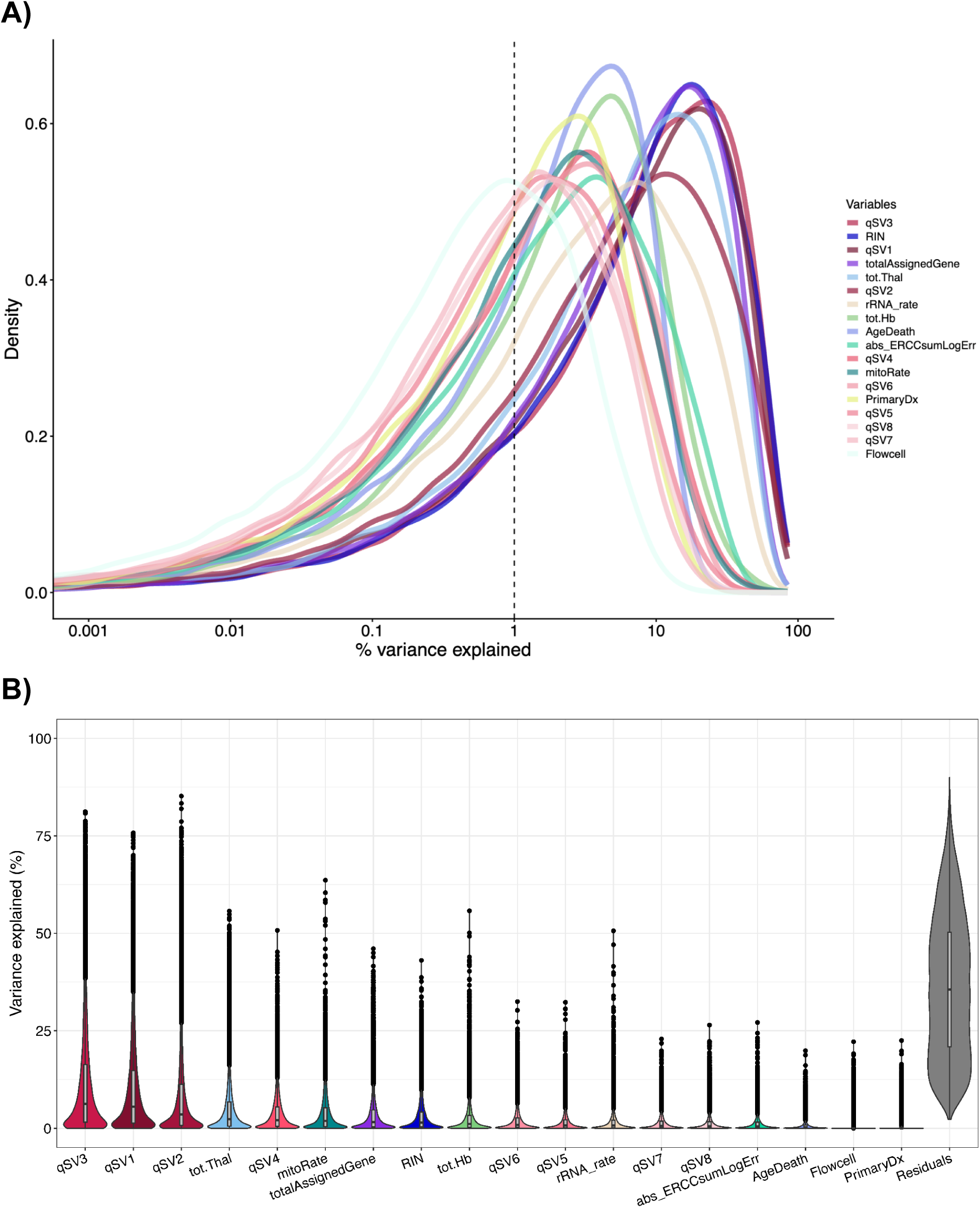
Bulk RNA-sequencing variance partition across covariates. **A**) Density lines for the percent of variance explained by each covariate across all genes, as estimated using `plotExplanatoryVariables()` from the *scater* package. The x-axis is displayed on a log_10_ scale. The RNA Integrity Number (RIN), quality Surrogate Variables (qSV) 1 and 3, percent of reads assigned to genes (totalAssignedGene), and total percent of Thalamus cell types estimated by deconvolution (tot.Thal) are among the covariates that explain the highest percent of variance. **B**) Variance explained distribution boxplots across all genes for each of the covariates, as estimated with the *variancePartition* package.

**Fig. S18.**
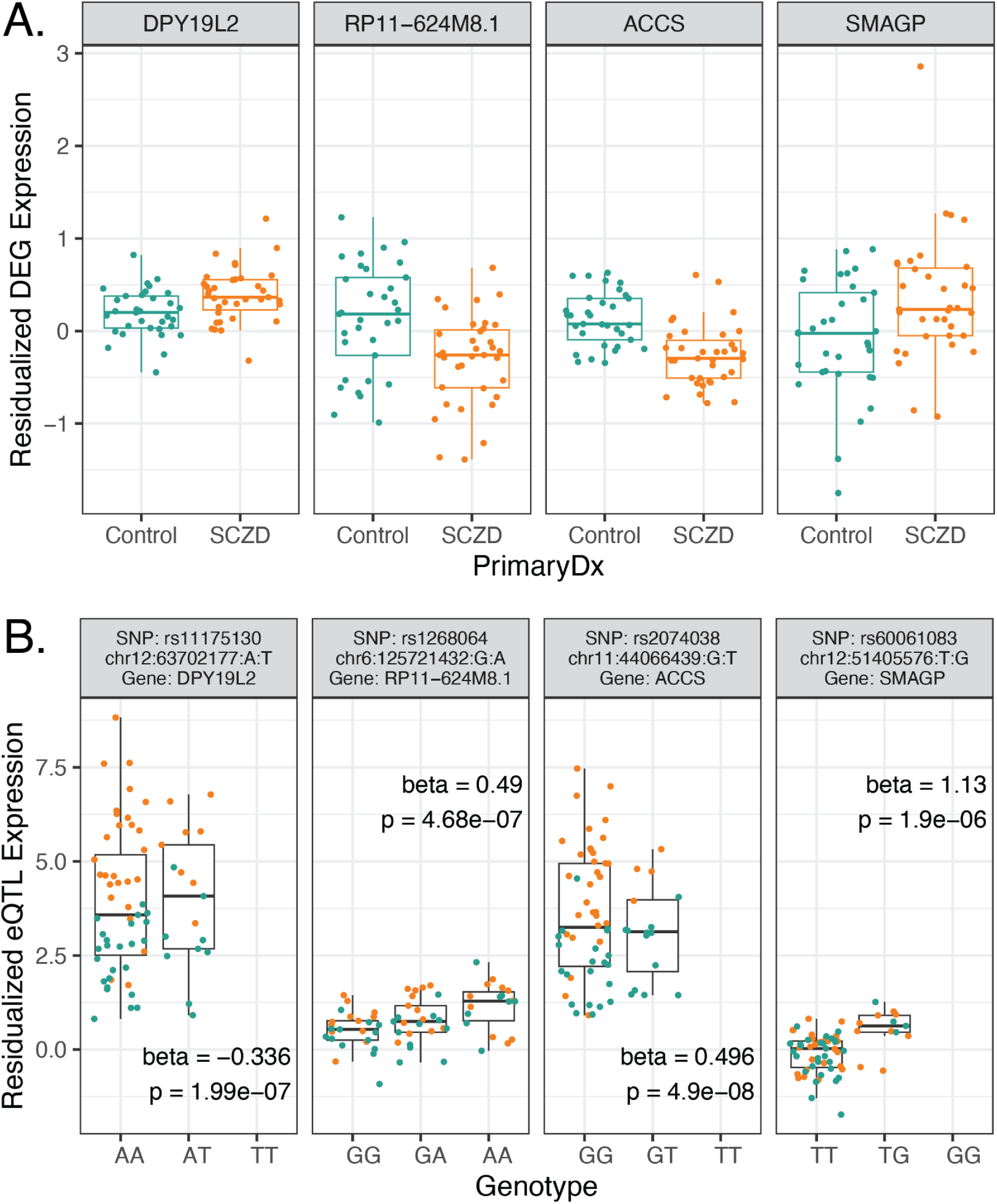
Residualized normalized expression by diagnosis and genotype for eQTLs involving SCZD DEGs. The remaining four of seven such eQTLs are shown here, with the first three in **Fig. 5A**. **A)** Residualized normalized expression by diagnosis for SCZD DEGs (FDR < 0.1) found in independent eQTLs (FDR < 0.05). **B)** Residualized normalized expression by genotype for SNPs paired with SCZD DEGs and constituting independent eQTLs, labeled with eQTL regression (beta) and nominal association p-values.

**Fig. S19.**
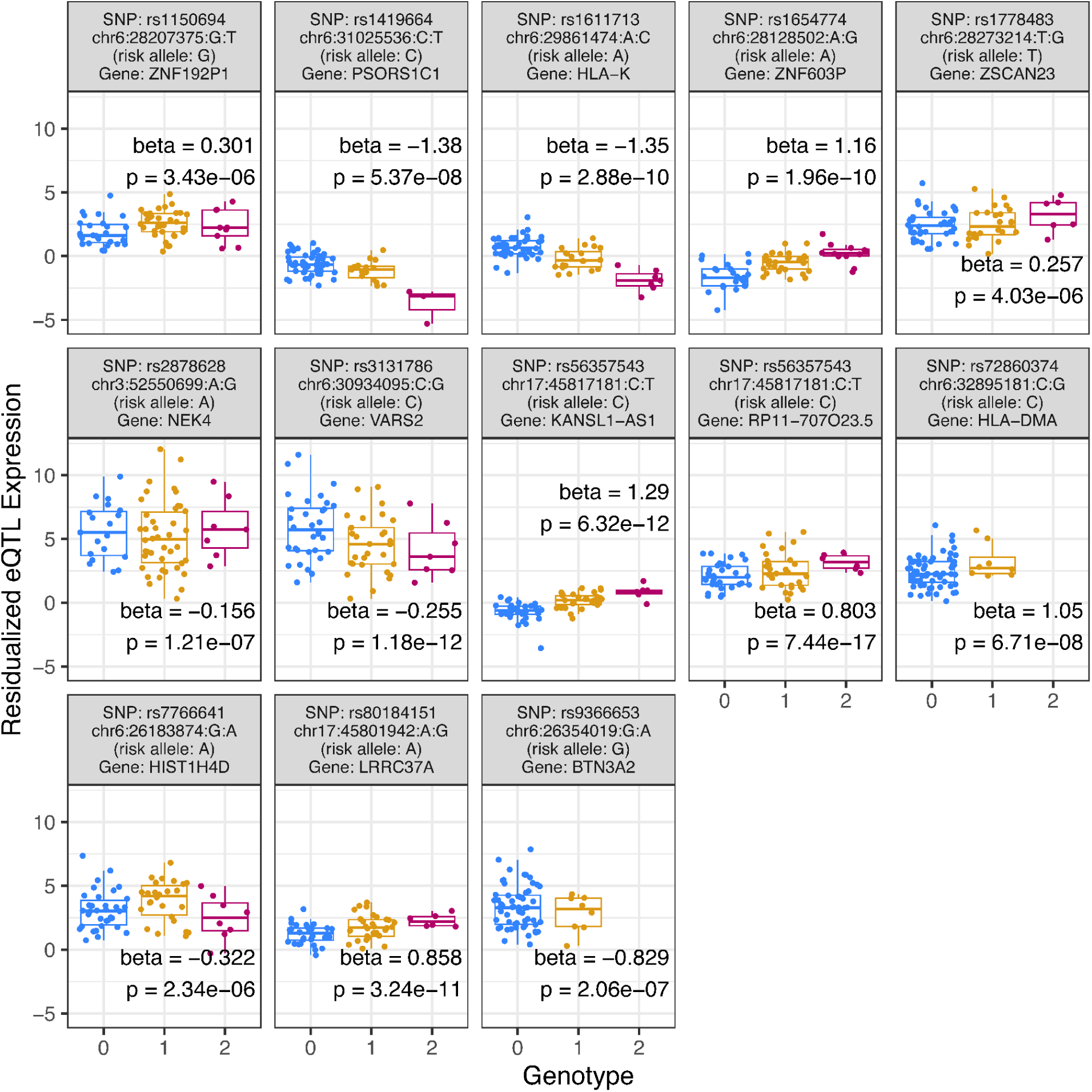
Independent eQTLs SNP-gene pairs with SNPs overlapping PGC3 SCZD GWAS risk SNPs. Boxplots by SNP alleles of residualized normalized expression for the remaining 13 genes from independent eQTLs whose SNPs are PGC3 SCZD GWAS risk SNPs (p<5 * 10^-8^). The other three are shown in **Fig. 5B**. Plots include eQTL regression (beta) and nominal association p-value. For the *X* axis, 0: homozygous major allele, 1: heterozygous, 2: homozygous minor allele.

**Fig. S20.**
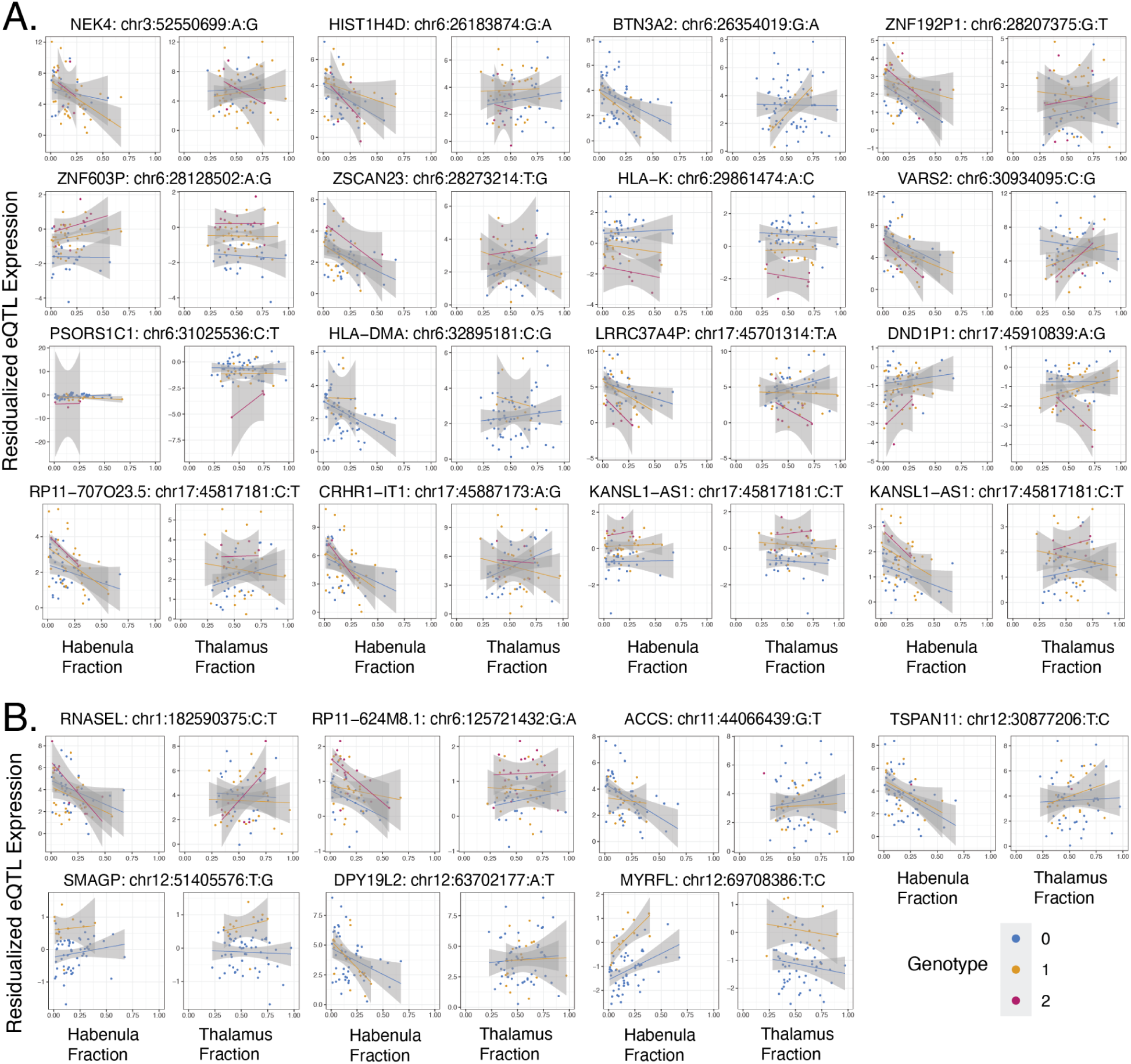
Habenula and thalamus neuronal cell fraction interaction with SNP alleles for independent eQTLs whose SNPs are SCZD risk SNPs or genes are DEGs. Scatterplots of residualized normalized expression compared against either the estimated habenula or the thalamus neuronal fractions (`Tot.Hb` and `Tot.Thal`) with SNP allele regression lines colored by allele (homozygous major allele in blue, heterozygous in orange, homozygous minor allele in red). In all SNP-gene pairs, the major allele matched the reference allele. A) Cell fraction interaction eQTLs could be computed for 14 of the 16 SNP-gene pairs from Hb-enriched thalamus independent eQTLs whose SNPs are PGC3 SCZD GWAS risk SNPs (p<5 * 10^-8^). B) SNP-gene pairs from Hb-enriched thalamus independent eQTLs whose genes were SCZD DEGs (FDR<0.1). Related to **Fig. 5C**.

## Supplementary Tables

**Table S1. Donor demographics and bulk RNA-sequencing *SPEAQeasy* metrics.** Donor ID (BrNum), bulk RNA-seq library ID (RNum), age at time of death, sex, race, and primary diagnosis (SCZD vs. Control) are the demographics included. *SPEAQeasy* metrics are documented at https://research.libd.org/SPEAQeasy/outputs.html#quality-metrics. The bulk RNA-seq sequencing flowcell, metrics derived from the bulk RNA-seq QC analysis with *scran,* deconvolution results with *BisqueRNA*, quality surrogate variables (qSVs) obtained with *qsvaR*, and the first 10 DNA genotype ancestry principal components (snpPCs) are also included. All snRNA-seq samples are a subset of these donors.

**Table S2. CellRanger snRNA-seq metrics.** Metrics computed by *CellRanger* v6.0.0 for the seven snRNA-seq samples. For more details about these metrics see: https://www.10xgenomics.com/support/software/cell-ranger/latest/analysis/outputs/cr-3p-outputs-metrics-count

**Table S3. Top 50 mean ratio marker genes by fine cell type category.** For each of the seventeen cell type categories – including the finer habenula cell type subclusters – the top 50 mean ratio marker genes are shown.

**Table S4. Donor demographics for RNAScope smFISH experiments.** Brain donor ID (BrNum), age at time of death (in years), sex, race, primary diagnosis, estimated postmortem interval (PMI, in hours), RNA integrity number (RIN) screened on the Prefrontal Cortex (PFC), and toxicology information (Positive at Time of Death) are provided for the three independent donors used in these experiments.

**Table S5. RNAScope experimental design summary.** Experimental design for RNAScope experiments. Probes, catalog numbers, channel and Opal dye assignments, and dye concentrations used for each probe combination.

**Table S6. Top 50 mean ratio marker genes by broad cell type category.** For each of the nine broad cell type categories, the top 50 mean ratio marker genes are shown. Only the top 25 mean ratio marker genes by cell type were used for estimating the cell type proportions by deconvolution.

**Table S7. Bulk RNA-seq Schizophrenia vs. Control differential gene expression (DGE) analysis results.** Harmonized gene-level results for the Habenula (Hb), dorsolateral Prefrontal Cortex (DLPFC) and Hippocampus (HIPPO) from BrainSEQ Phase II, Caudate from BrainSEQ Phase III, and Dentate Gyrus (DG). This analysis was restricted to genes expressed in the Hb.

**Table S8. Independent Hb-enriched eQTL SNP-gene pairs.** *tensorQTL* outputs are provided for the 717 significant (FDR < 0.05) independent eQTLs. Available interaction eQTL statistics with `Tot.Hb` and `Tot.Thal` fractions are provided for the same eQTLs. Related to **Fig. 5**.

**Table S9. Gene eQTL vs SCZD DEG, and SNP eQTL vs SNP SCZD risk overlap summary.** Two by two tables comparing whether **A)** a gene is a part of a SNP-gene eQTL pair or an SCZD DEG, or **B)** whether a SNP is part of an SNP-gene eQTL pair or an SCZD PGC3 GWAS risk SNP. These tables are the input for computing Fisher’s exact test for a positive association. SNPs were subset to those present in both the Hb-enriched data as well as the PGC3 GWAS data. Related to **Fig. 5**.

**Table S10. eQTL and SCZD PGC3 colocalization summary results.** Results extracted from `coloc::coloc.abf()$results` for all SNPs in the 95% credible set for H_4_ in each of the genes that had a posterior probability of H_4_ greater than 0.8. In addition to the columns computed by `coloc.abf()`, we included the gene ID and symbol on which the ±500kb window is centered, the 1Mb window summary `PP.H4.abf` value (filtered to genes with `SNP.PP.H4` > 0.8), whether the gene was identified as a SCZD DEG (FDR<0.1), whether the SNP is a SCZD PGC3 risk SNP (p<5*10^-8^), or whether the colocalized SNP-gene pair is also an independent eQTL SNP-gene pair (FDR<0.05).

## Notes

https://github.com/LieberInstitute/Habenula_Pilot

### Summary of Updates

This revised version includes new results that led to a new main figure (Figure 5), and 3 new supplementary tables, and 3 new supplementary figures. These new results are from eQTL and GWAS colocalization analyses.

